# *Bacillus subtilis* YngB contributes to wall teichoic acid glucosylation and glycolipid formation during anaerobic growth

**DOI:** 10.1101/2020.11.30.405621

**Authors:** Chih-Hung Wu, Jeanine Rismondo, Rhodri M. L. Morgan, Yang Shen, Martin J. Loessner, Gerald Larrouy-Maumus, Paul S. Freemont, Angelika Gründling

## Abstract

UTP-glucose-1-phosphate uridylyltransferases (UGPases) are enzymes that produce UDP-glucose from UTP and glucose-1-phosphate. In *Bacillus subtilis* 168, UDP-glucose is required for the decoration of wall teichoic acid (WTA) with glucose residues and the formation of glucolipids. The *B. subtilis* UGPase GtaB is essential for UDP-glucose production under standard aerobic growth conditions, and *gtaB* mutants display severe growth and morphological defects. However, bioinformatics predictions indicate that two other UGPases, are present in *B. subtilis*. Here, we investigated the function of one of them named YngB. The crystal structure of YngB revealed that the protein has the typical fold and all necessary active site features of a functional UGPase. Furthermore, UGPase activity could be demonstrated *in vitro* using UTP and glucose-1-phosphate as substrates. Expression of YngB from a synthetic promoter in a *B. subtilis gtaB* mutant resulted in the reintroduction of glucose residues on WTA and production of glycolipids, demonstrating that the enzyme can function as UGPase *in vivo*. When wild-type and mutant *B. subtilis* strains were grown under anaerobic conditions, YngB-dependent glycolipid production and glucose decorations on WTA could be detected, revealing that YngB is expressed from its native promoter under anaerobic condition. Based on these findings, along with the structure of the operon containing *yngB* and the transcription factor thought to be required for its expression, we propose that besides WTA, potentially other cell wall components might be decorated with glucose residues during oxygen limited growth condition.

## Introduction

The cell envelope of bacteria is composed of several sugar-containing polymers, including peptidoglycan, capsular polysaccharides, lipopolysaccharide (LPS) in Gram-negative bacteria and secondary cell-wall polymers such as teichoic acids in Gram-positive bacteria (1–5). Secondary cell-wall polymers in Gram-positive bacteria can either be complex and made up of different repeating sugar units, or more simple glycerol- or ribitol-phosphate polymers that are further decorated with sugar residues (6–8). Under standard aerobic growth conditions, the model Gram-positive organism *Bacillus subtilis* strain 168 produces two different types of teichoic acid (TA). Lipoteichoic acid (LTA) is a polyglycerolphosphate polymer that is linked by the glycolipid anchor diglucosyl-diacylglycerol (Glc_2_-DAG) to the outside of the bacterial membrane and further decorated with D-alanine and N-acetylglucosamine (GlcNAc) residues, and wall teichoic acid (WTA) is a polyglycerol phosphate polymer covalently linked to peptidoglycan and decorated with D-alanine and glucose residues (9,10). Both polymers are made up of glycerol phosphate repeating units but are produced by separate pathways. Whereas LTA is polymerized on the outside of the cell, WTA is polymerized within the cell (7,8). Under phosphate-limiting growth conditions, at least part of the WTA is replaced with teichuronic acid, a non-phosphate containing anionic cell-wall polymer (11,12).

For the synthesis as well as decoration of bacterial cell-wall polymers, several important enzymes producing nucleotide-activated sugars are required (13,14). For the decoration of LTA with GlcNAc residues, UDP-GlcNAc is utilized, which is also an essential precursor for peptidoglycan and WTA synthesis (7,15). UDP-GlcNAc is produced from glucosamine-1-phosphate by the bifunctional enzyme GlmU, which has acyltransferase and uridylyltransferase activity (16). The nucleotide-activated sugar is subsequently used by a multi-enzyme glycosylation machinery for the modification of LTA with GlcNAc residues on the outside of the cell. For this process, it is thought that the membrane-linked glycosyltransferase CsbB transfers the GlcNAc residue from UDP-GlcNAc onto the lipid carrier undecaprenyl phosphate (C_55_-P) to generate the lipid-linked sugar intermediate C_55_-P-GlcNAc (17–19). Next, this intermediate is transferred across the membrane with the aid of the small membrane protein and proposed flippase enzyme GtcA, which belongs to the GtrA protein family (20,21). The GlcNAc residues are finally added to the LTA polymer by the multi-membrane spanning GT-C-type glycosyltransferase YfhO (19).

The glycosylation process of WTA in *B. subtilis* is much simpler. The glucose residues are attached to the polymer within the cytoplasm of the cell by the glycosyltransferase TagE using UDP-glucose as substrate (22,23). The nucleotide activated sugar precursor UDP-glucose is produced from UTP and glucose-1-phosphate by UTP-glucose-1-phosphate uridylyltransferase (UGPase) enzymes (14). UGPases are widespread in bacteria and are often named GalU. For several Gram-positive as well as Gram-negative bacterial pathogens, GalU has been shown to be required for full virulence due to its involvement in biofilm formation, capsule and/or LPS biosynthesis (24–27). A well-characterized GalU equivalent in *B. subtilis* is GtaB, and the first *gtaB* mutants were isolated as part of studies investigating phage-resistant *B. subtilis* strains (10). These studies also revealed that the phage resistance is due to the lack of glucose decorations on WTA (28,29). In *B. subtilis* 168, UDP-glucose is not only required for the decoration of WTA with glucose residues but also for the production of glycolipids. An abundant glycolipid found in the membrane is Glc_2_-DAG, which serves as the lipid anchor for LTA and is produced by the transfer of two glucose molecules from UDP-glucose onto the membrane lipid diacylglycerol (DAG) by the glycosyltransferase UgtP (or sometimes also named YpfP) (30–33). Several independently obtained *B. subtilis gtaB* mutants have now been characterized. These mutants are resistant to certain types of phages, display morphological and growth defects, lack glucose decorations on WTA and are unable to produce glycolipids under standard laboratory growth conditions (31,34). Furthermore, no UGPase activity could be detected in lysates prepared from *gtaB* mutant strains, indicating that GtaB is the sole functional UGPase in *B. subtilis* under these conditions (28). However, apart from GtaB, two other predicted UGPases, YtdA and YngB are encoded in the *B. subtilis* 168 genome: as part of this study we investigated the function of YngB (35,36).

The *yngB* gene is thought to be part of the *yngABC* operon (35), but an additional internal promoter appears to be present in *yngB* driving *yngC* expression. Furthermore, expression of *yngC* has been reported to be under control of a sigma M-dependent promoter (37). YngA belongs to the GtrA protein family and could therefore, similar as proposed for the GtcA enzyme involved in the LTA glycosylation process, function as flippase enzyme and be required for the transfer of a sugar-linked C_55_-P lipid intermediate across the membrane. YngC is a membrane protein belonging to the DedA family of proteins. The function of the protein is not clear, but proteins belonging to this family have been shown to be important for multiple processes in other bacteria, including cell division, membrane composition and antibiotic resistance (38–40). The transcription of the *yngABC* operon is activated by the transcription factor YclJ, which forms a two-component system (TCS) with the histidine sensor kinase YclK (35,41,42). The biological role of the YclJ-YclK TCS remains unknown but it has been shown that the *yclJK* regulon is upregulated during low oxygen conditions (43,44). Hence the proposed UGPase YngB could potentially function under oxygen-limiting conditions. However, the expression might be more complex as the expression of the *yclJK* genes in turn is controlled by the ResDE two-component system, a signal transduction system known to play a key role in the expression of both aerobic and anaerobic respiration-related genes in *B. subtilis* (45,46).

In this study, we aimed to determine if YngB is a functional UGPase enzyme and to provide insight into its biological function. Based on its structure as well as its *in vitro* enzymatic activity, we showed that YngB is a functional UGPase. YngB can also function *in vivo* as UGPase leading to the decoration of WTA with glucose residues and glycolipid production in the absence of GtaB, when expressed from a synthetic promoter. YngB-dependent glycosylation of WTA and glycolipid production was also observed when *B. subtilis* was grown under anaerobic fermentative growth condition. This revealed that besides GtaB, YngB is expressed from its native promoter under anaerobic growth conditions and functions as UGPase. Based on these findings and previous reports on the transcription control of the *yngABC* and other operons in *B. subtilis*, the potential decoration of other cell wall structures with glucose residues during oxygen limited growth condition will be discussed.

## Results

### The crystal structure of the *B. subtilis* YngB protein reveals a putative UDP-glucose binding site

Three paralogous UGPases, GtaB (BSU_35670), YtdA (BSU_30850) and YngB (BSU_18180) are encoded in the genome of *B. subtilis* strain 168 (Fig. S1). GtaB has been characterized as a UGPase and in its absence *B. subtilis* lacks glucose decorations on WTA and is unable to produce glycolipids during vegetative growth (28,31). Here we set out to determine whether YngB is a bona fide UGPase and to investigate its biological function. An amino acid sequence alignment of the *B. subtilis* proteins GtaB, YngB and YtdA with the UGPase enzymes A4JT02 from *Bulkholderia vietnamiensis* (PDB code: 5i1f), GalU_Hp_ from *Helicobacter pylori* (PDB code: 3juk) and GalU_Cg_ from *Corynebacterium glutamicum* (PDB code: 2pa4), for which structures with bound UDP-glucose are available (47,48), revealed that most of the UDP-glucose binding residues (Fig. S1, colored in yellow) as well as the metal-chelating residue (Fig. S1, colored in cyan) are conserved in the three *B. subtilis* proteins (14,47,48). To determine the crystal structure of YngB, selenomethionine-substituted protein was produced in *E. coli* and purified as a C-terminal His-tag fusion protein. The protein crystallized as a dimer in the asymmetric unit and the structure was solved at 2.80 Å by experimental phasing (Fig.1A and Table 1). YngB displayed a Rossmann fold with alternating α-helices and β-strands, which is commonly found in nucleotide-binding proteins (49). More specifically, the YngB monomer contains a central β-sheet surrounded by α-helices (47,50). Similar to the dimer interactions described for homologous UGPases, hydrogen bonds are formed between Tyr102 residues on the two β-strands located at the interface of the monomers, producing an extended central β-sheet spanning across the two subunits (50,51). The overall structure of YngB is very similar to homologous UGPases including A4JT02, GalU_Hp_ and GalU_Cg_ as shown by superimposition (Fig. S2). The outer most C-terminal α-helix is one of the most variable region and either absent in GalU_Hp_ or in a different conformation in GalU_Cg_ (Fig. S2). To locate the substrate-binding site, the YngB and UDP-glucose bound *H. pylori* GalU_Hp_ structures (PDB code: 3juk) (47) were overlayed. The structures superimposed with an r.m.s.d. of 1.61 Å and the putative UDP-glucose binding pocket could easily be identified in YngB (Fig.1B & 1C). Based on this alignment it can be predicted that residues Gly110, Gln105, Ala13, Gly14, Glu29, and Lys28 in YngB interact with the uridine moiety; Asp133 is involved in the chelation of a Mg^2+^ ion; Asp134, Val204, Gly172, and Glu191 interact with the glucose moiety; and Lys192 interacts with the diphosphate moiety (Fig. 1C). Based on this structural analysis, YngB has all the required features expected of a bona fide UGPase enzyme.

**Table 1.** Crystallographic data and refinement statistics

|  | **YngB (SeMet)** |
| --- | --- |
| **Data collection** |  |
| Space group | I 21 21 21 |
| Cell dimensions |  |
| a, b, c (Å) | 53.91 158.61 179.50 |
| $\alpha$, $\beta$, $\gamma$ (°) | 90.000 90.000 90.00 |
| Wavelength (Å) | 0.9794 |
| Resolution (Å) | 51.09-2.80(2.95-2.80) |
| No. reflections | 36539(5279) |
| Unique reflections | 19457(2777) |
| R<sub>merge</sub> | 0.067(0.468) |
| R<sub>meas</sub> | 0.095(0.661) |
| CC<sub>(1/2)</sub> | 0.994(0.695) |
| I / $\sigma(I)$ | 5.6(1.2) |
| Completeness (%) | 99.8(98.8) |
| Redundancy | 1.9(1.9) |
| Wilson B-factor (Å<sup>2</sup>) | 79.0 |
| No. of protein molecules/ asym. unit | 2 |
| **Refinement** |  |
| Resolution (Å) | 51.09-2.80 |
| No. reflections all/free | 19535/937 |
| Rwork / Rfree | 0.191/0.244 |
| No. of atoms | A 2197; B 2181 |
|  | Water 54 |
| B factors (Å<sup>2</sup>) |  |
| Protein | A 81.9; B 89.4 |
| Water | 72.58 |
| Clashscore | 19 |
| R.m.s. deviations |  |
| Bond lengths (Å) | 0.0070 |
| Bond angles (°) | 1.792 |
| Ramachandran plot |  |
| Favored (%) | 89 |
| Allowed (%) | 11 |
| Outliers (%) | 0 |
| PDB accession code | 7B1R |

Statistics in brackets refer to the highest resolution shell

**Figure 1.**
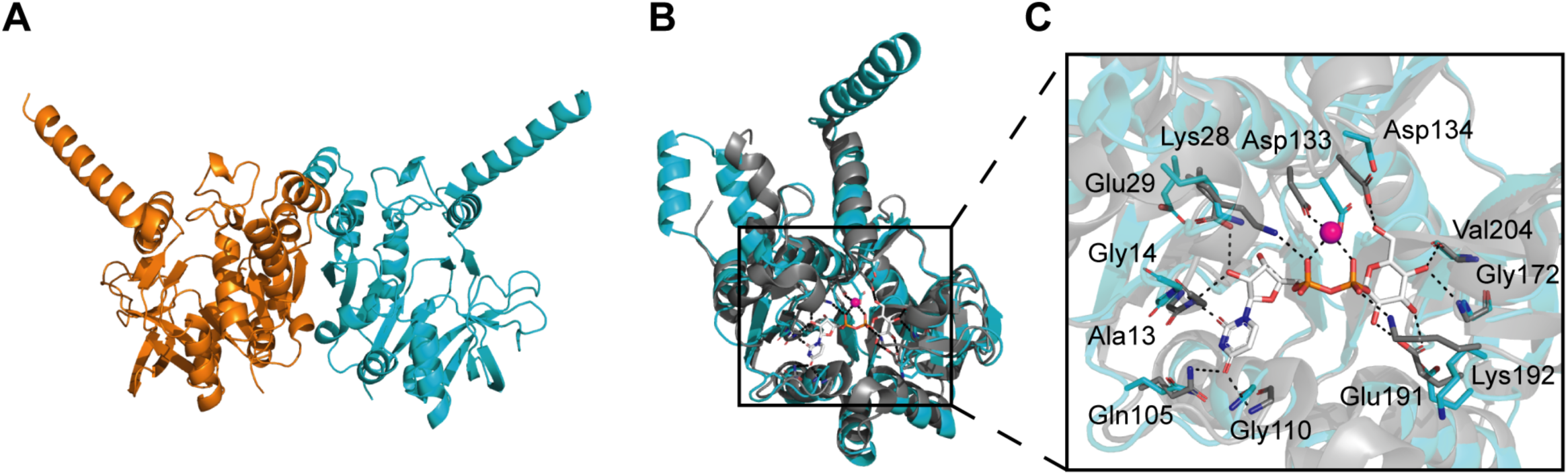
YngB crystal structure and proposed UDP-glucose binding site residues. *A*, crystal structure of YngB shown in cartoon representation. The protein crystalized as a dimer in the asymmetric unit, and individual monomers are shown in orange and cyan respectively. *B*, overlay of the *B. subtilis* YngB (cyan) and the UDP-glucose bound *H. pylori* GalU_HP_ (3JUK) (grey) structures. The structures are shown in ribbon representation, with UDP-glucose and UDP-glucose binding residues shown in stick representation. *C*, enlarged view of the substrate binding site from the alignment shown in panel B. The magenta sphere represents a magnesium ion; ionic interactions with the magnesium ion and hydrogen bonds are shown as dashed lines. Proposed UDP-glucose binding residues in YngB are Gly110, Gln105, Ala13, Gly14, Glu29, and Lys28 interacting with the uridine moiety; Asp133 chelating the magnesium ion; Asp134, Val 204, Gly172, and Glu191 interacting with the glucose moiety; and Lys 192 interacting with the diphosphate moiety.

### YngB shows UGPase activity *in vitro*

To determine if YngB has UGPase activity *in vitro*, enzyme assays were preformed using a method previously described for assessing the UGPase activity of the GalU enzyme from *Erwinia amylovora* (52). For this assay, purified proteins are incubated with α-glucose-1-phosphate (G-1-P) and UTP and active UGPases will convert these substrates into UDP-glucose and pyrophosphate. The generated pyrophosphate is then hydrolyzed by a pyrophosphatase to two molecules of inorganic phosphate, which is quantified calorimetrically. To assess the enzymatic activity of YngB, assays were performed with recombinant YngB protein using recombinant GtaB protein as control, and increasing concentrations of G-1-P and a fixed concentration of UTP or increasing concentrations of UTP and fixed concentration of G-1-P. These experiments revealed that YngB possesses UGPase activity, and Michaelis-Menten curves could be produced for both enzymes acting on either substrate (Fig. 2). From these data, apparent K_m_ values of 45.6 ± 3.24 µM (GtaB) and 42.1 ± 20.2 µM (YngB) were calculated for G-1-P and 49.5 ± 10.2 µM (GtaB) and 62.9 ± 13.8 µM (YngB) for UTP, respectively. k_cat_ values of 1.06 s^-1^ (GtaB) and 0.264 s^-1^ (YngB) were calculated for G-1-P and 1.04 s^-1^ (GtaB) and 0.293 s^-1^ (YngB) for UTP, respectively. These enzyme assays confirmed that YngB has UGPase activity *in vitro* and revealed that under the assay conditions used, YngB had a lower turnover number compared to GtaB.

**Figure 2.**
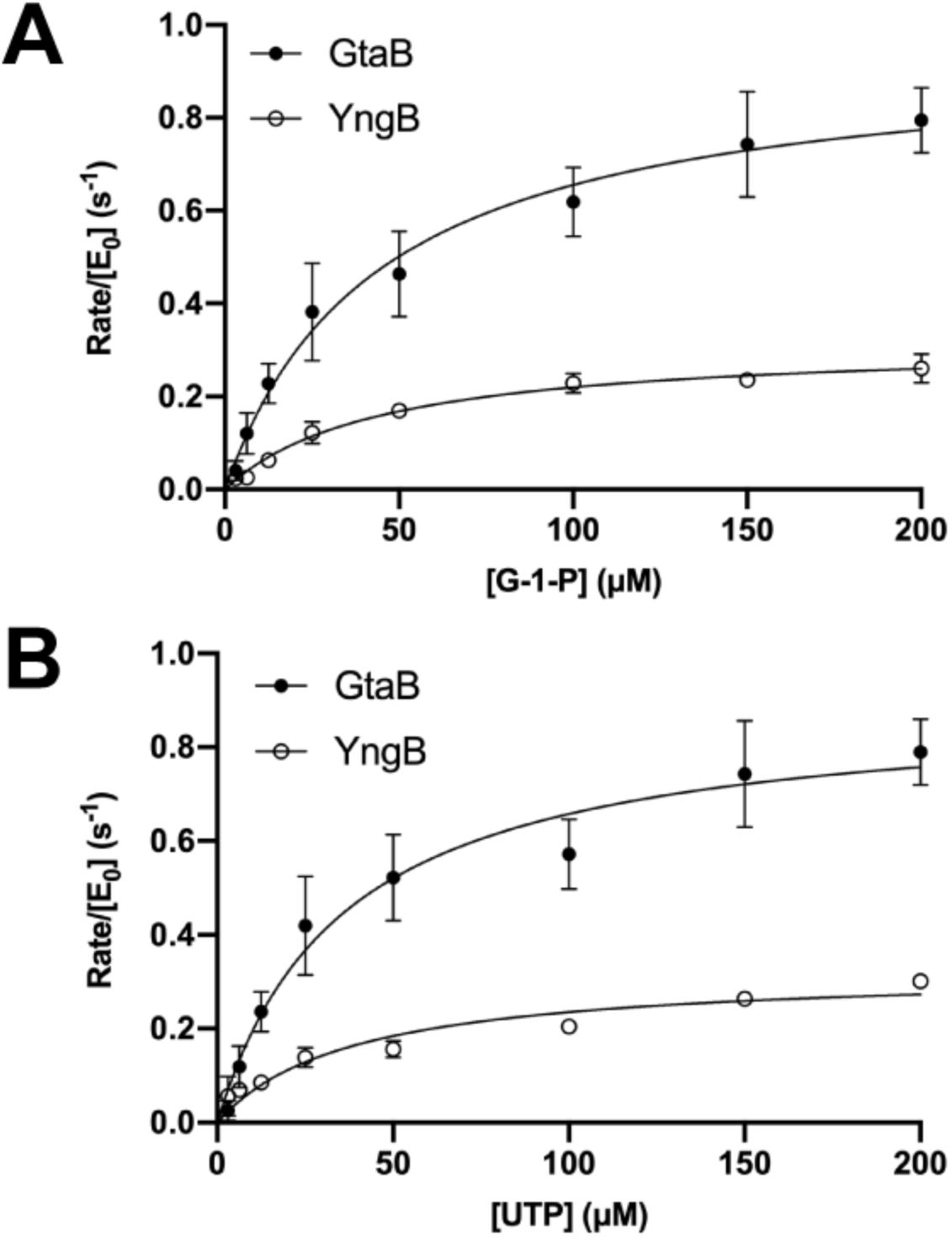
Enzyme activity of GtaB and YngB. (*A-B*), Michaelis-Menten kinetics using purified GtaB or YngB proteins and *A*, increasing concentrations of the substrate glucose-1-phosphate (G-1-P) or *B*, increasing concentrations of the substrate UTP. The number of UDP-glucose formed per molecule of GtaB or YngB per second is plotted versus substrate concentration. The measured apparent K_m_ values for glucose-1-phosphate were 45.6 ± 3.24 µM (GtaB) and 42.1 ± 20.2 µM (YngB) and for UTP, 49.5 ± 10.2 µM (GtaB) and 62.9 ± 13.8 µM (YngB), respectively. The experiment was performed three times with technical replicates. A representative graph from one experiment and plotting the mean and SD from the technical replicates is shown. The SD values for some data points were too small to be displayed on the graph. Michaelis-Menten curves were produced with Prism and K_m_ values given are the mean ± SD from the three independent experiments.

### Expression of YngB from an inducible promoter leads to glucosylation of WTA

In *B. subtilis* strain 168, glucose is transferred onto WTA by the glycosyltransferase TagE using UDP-glucose as substrate (23). Under standard aerobic growth conditions, GtaB appears to be the only enzyme that produces UDP-glucose, as glucose is absent from WTA in *gtaB* mutant strains (28,34,53). This is somewhat at odds with our data showing that *B. subtilis* YngB protein is a functional UGPase enzyme. Possible explanations could be that YngB uses a different sugar or nucleotide as substrate *in vivo* or that it is not expressed under standard aerobic growth conditions. To address these issues, the *yngB* gene or, as a control, the *gtaB* gene, were placed under control of the synthetic IPTG-inducible P_hyperspank_ promoter (short P_hyper_ promoter) and introduced into the chromosome of the Δ*gtaB* single mutant and the Δ*gtaB*Δ*yngB* double mutant strain. The presence of glucose on WTA in the different *B. subtilis* strains was initially assessed by fluorescence microscopy after staining the bacteria with fluorescently labeled concanavalin A, a lectin that specifically binds to terminal glucose residues (34). The different *B. subtilis* strains were grown aerobically in medium supplemented with IPTG and culture samples were taken at mid-log growth phase for microscopy analysis. A fluorescence signal was observed for the wild-type and Δ*yngB* single mutant strain, but as expected was absent from the Δ*gtaB* single and the Δ*gtaB*Δ*yngB* double mutant strains (Fig. 3). Consistent with previous observations, Δ*gtaB* mutant bacteria displayed morphological defects and the cells were curled and showed some bulges, which was also seen for Δ*gtaB*Δ*yngB* double mutant cells (Fig 3). Expression of either *yngB* or *gtaB* from the inducible promoter (P_hyper_) in the Δ*gtaB* single or Δ*gtaB*Δ*yngB* double mutant strains fully or at least partially complemented both phenotypes; the cells showed again binding to the fluorescent lectin and bacteria complemented with GtaB had a normal rod-shaped morphology and the bacteria complemented with YngB showed an improved cell morphology and the cell were less curved (Fig. 3). These results indicate that YngB is not expressed from its native promoter under standard aerobic growth conditions; however, when expressed from an inducible promoter, YngB can function as UGPase *in vivo*. To further confirm that YngB expression leads to the decoration of WTA with glucose resides, WTA was isolated from wild-type, the Δ*gtaB* mutant and strains Δ*gtaB* P_hyper_-*gtaB* and Δ*gtaB* P_hyper_-*yngB* and analyzed by NMR and LC-MS. The ^1^H NMR spectrum obtained for WTA isolated from the wild-type strain revealed a peak at 5.2 ppm likely derived from the hydrogen atom at the anomeric carbon of the glucose residue on WTA (23) (Fig. 4A). This peak was absent in the sample derived from the Δ*gtaB* mutant strain but could again be detected in samples isolated from strains Δ*gtaB* P_hyper_-*gtaB* and Δ*gtaB* P_hyper_-*yngB* (Fig. 4A), indicating that WTA is indeed decorated with glucose residues upon expression of YngB from an inducible promoter. For the LC-MS analysis, the purified WTA was hydrolyzed with hydrogen fluoride and the depolymerized species were characterized. A species with a mass of 253.09 (*m/z*) corresponding to a glycerol-glucose repeating unit was detected for samples derived from the wild-type as well as strains Δ*gtaB* P_hyper_-*gtaB* and Δ*gtaB* P_hyper_-*yngB* (Fig. 4B) but was absent from the Δ*gtaB* sample (Fig. 4B). These data highlight that YngB can function as UGPase *in vivo* and produce UDP-glucose, which can subsequently be used for the decoration of WTA with glucose moieties.

**Figure 3.**
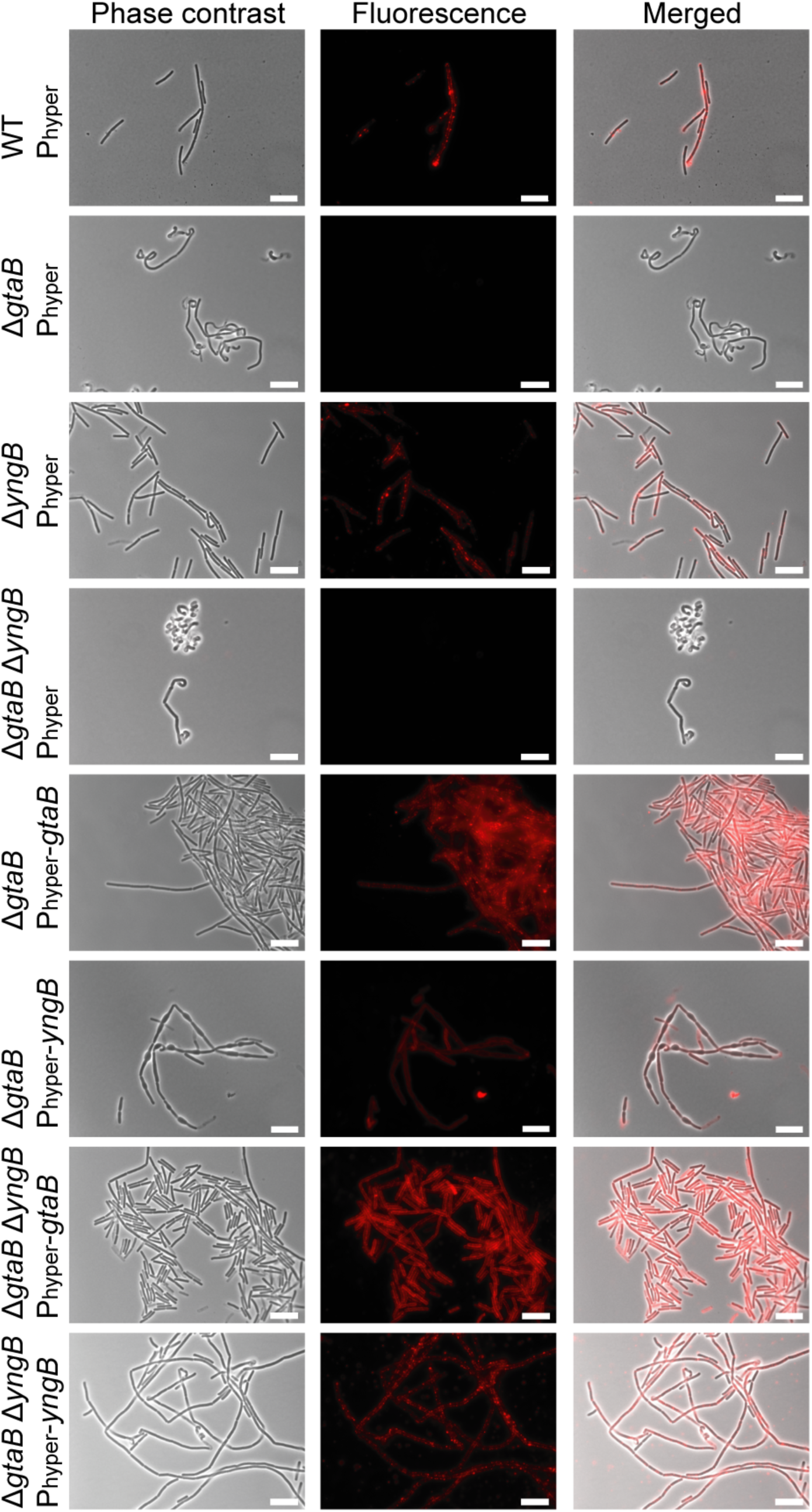
Microscopy analysis and detection of glucose modifications on WTA produced by aerobically grown wild-type *B. subtilis*, mutant and complementation strains. The indicated *B. subtilis* strains were grown under aerobic growth conditions to mid-log phase in LB medium supplemented with IPTG and glucose. Bacteria were prepared for microscopy analysis and stained with the fluorescently labelled Alexa Fluor^™^ 594 Concanavalin A lectin to detect glucose modifications on WTA as described in the experimental procedure section. The experiment was performed 3 times and representative phase contrast, fluorescence and merged images are shown for each strain. Scale bars represent 10 µm.

**Figure 4.**
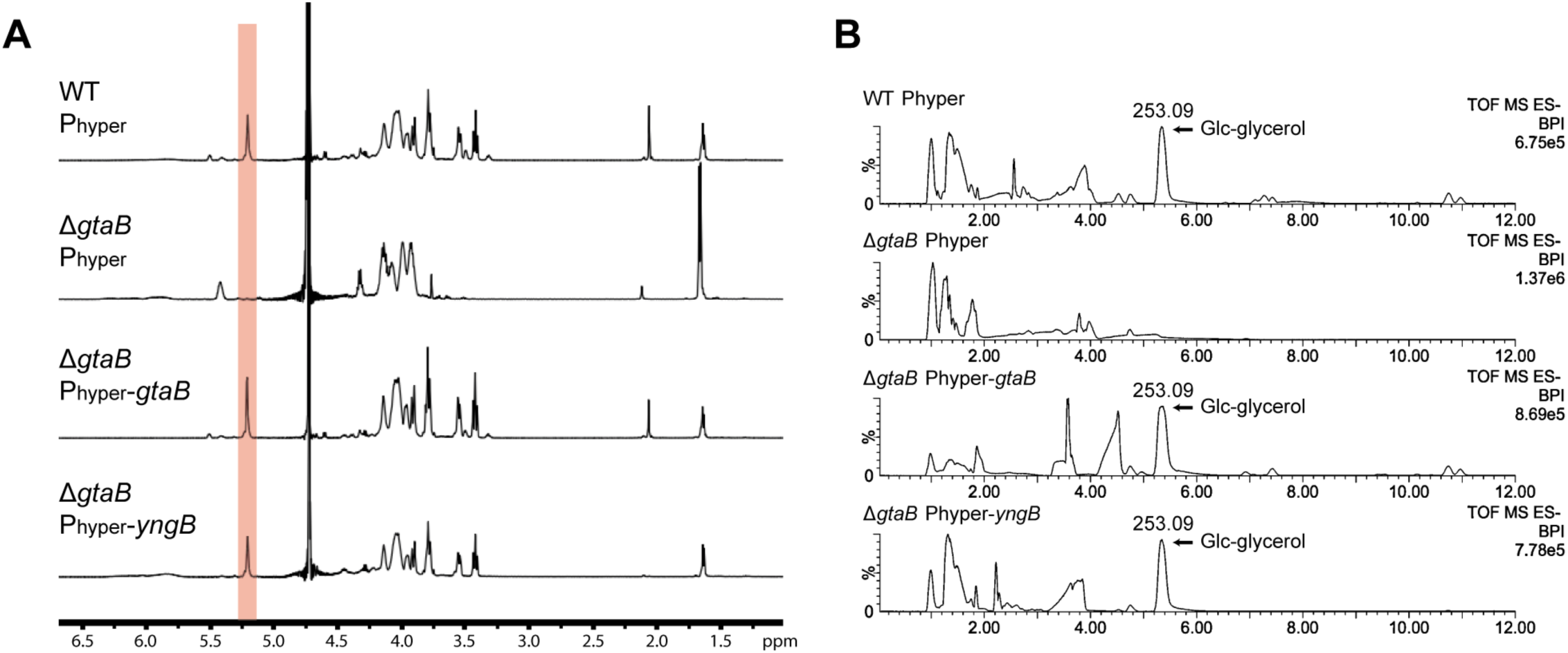
NMR and UPLC-MS analysis of WTA isolated from wild-type *B. subtilis*, mutant and complementation strains. *A*, ^1^H NMR spectra of WTA isolated from wild-type *B. subtilis* (WT P_hyper_), mutant Δ*gtaB* P_hyper_, and complementation strains Δ*gtaB* P_hyper_-*gtaB* and Δ*gtaB* P_hyper_-*yngB*. The peak at 5.2 ppm (highlighted in orange) is likely derived from the hydrogen atom at the anomeric carbon of glucose residues on WTA (23). The experiment was performed twice, and similar results were obtained with the peak at 5.2 ppm absent in the *gtaB* mutant but present in all other strains. The spectra from one experiment are shown. *B*, UPLC-MS analysis of hydrolyzed WTA samples derived from the same strains as described in *A*. The fragment with a mass signal of 253.09 (*m*/*z*) corresponding to glucose-glycerol species is absent in the *gtaB* mutant but present in all other strains.

### Expression of YngB from an inducible promoter leads to the formation of glycolipids

UDP-glucose is also used for the production of glycolipids in *B. subtilis* and the glycosyltransferase UgtP transfers one or more glucose moieties onto the membrane lipid diacylglyerol (DAG) (30,32). It has been reported that glycolipids are absent in *B. subtilis gtaB* mutant strains (31,54), however, our data suggest that expression of YngB in *gtaB* mutant strains should restore glycolipid production. To experimentally verify this, total membrane lipids were isolated from wild-type, Δ*gtaB* single and Δ*gtaB*Δ*yngB* double mutant strains as well as complementation strains expressing either *gtaB* (P_hyper_-*gtaB)* or *yngB* (P_hyper_-*yngB*) from the IPTG-inducible promoter. The isolated membrane lipids were separated by thin layer chromatography and glycolipids were visualized by staining with α-naphthol and sulfuric acid (Fig. 5). Several major bands were observed for the wild-type strain, two of which (Fig. 5 top and middle band) could be further characterized by MALDI-TOF MS (Fig.6) and MALDI-TOF MS/MS (Fig. S3). Based on the mass and fragmentation pattern, they are most consistent with Glc_2_-DAG (top band) and Glc_3_-DAG (middle band) sodium adducts with different fatty acid chain lengths (Table 2 and Table S1). These bands were absent in the Δ*gtaB* single and Δ*gtaB*Δ*yngB* double mutant strains, but again present upon expression of either *gtaB* or *yngB* from the IPTG-inducible promoter (Fig. 5A and 5B). The identity of these lipids was again consistent with sodium adducts of the glycolipids Glc_2_-DAG (top band) and Glc_3_-DAG (middle band) as assessed by MALDI-TOF MS and MALDI-TOF MS/MS (Table 2 and Table S1). The glycolipids produced by a Δ*yngB* mutant and complementation strain Δ*yngB* P_hyper_-*yngB* were also analyzed, however, no clear differences in the glycolipid profile were observed for these strains as compared to the wild-type strain (Fig. 5C). These data confirm that YngB is not required for glycolipid production under standard aerobic growth, presumably because the enzyme is not produced under these conditions and that GtaB is the main enzyme responsible for UDP-glucose production. However, when YngB is expressed from a synthetic promoter, it can take over the function of GtaB.

**Table 2.** Predicted and observed masses of glucolipid sodium adducts analyzed by MALDI-TOF mass spectrometry in the positive ion mode

| Possible fatty acid chain length | Chemical formula | Predicted molecular mass | Observed mass |  |  |
| --- | --- | --- | --- | --- | --- |
| | | | Wild-type | $\Delta gtaB$ P<sub>hyper-*gtaB*</sub> | $\Delta gtaB$ P<sub>hyper-*yngB*</sub> |
| Top band:<br>Glc<sub>2</sub>-DAG |  |  |  |  |  |
| (30:0) | C<sub>45</sub>H<sub>84</sub>Na<sub>1</sub>O<sub>15</sub> | 887.57 | 887.51 | 887.94 | 887.90 |
| (31:0) | C<sub>46</sub>H<sub>86</sub>Na<sub>1</sub>O<sub>15</sub> | 901.59 | 901.95 | 901.97 | 901.81 |
| (32:0) | C<sub>47</sub>H<sub>88</sub>Na<sub>1</sub>O<sub>15</sub> | 915.60 | 915.95 | 915.87 | 915.91 |
| Mid band:<br>Glc<sub>3</sub>-DAG |  |  |  |  |  |
| (30:0) | C<sub>51</sub>H<sub>94</sub>Na<sub>1</sub>O<sub>20</sub> | 1049.62 | 1049.85 | 1049.90 | 1049.77 |
| (31:0) | C<sub>52</sub>H<sub>96</sub>Na<sub>1</sub>O<sub>20</sub> | 1063.64 | 1063.85 | 1063.90 | 1063.93 |
| (32:0) | C<sub>53</sub>H<sub>98</sub>Na<sub>1</sub>O<sub>20</sub> | 1077.65 | 1077.85 | 1077.38 | 1077.90 |

**Figure 5.**
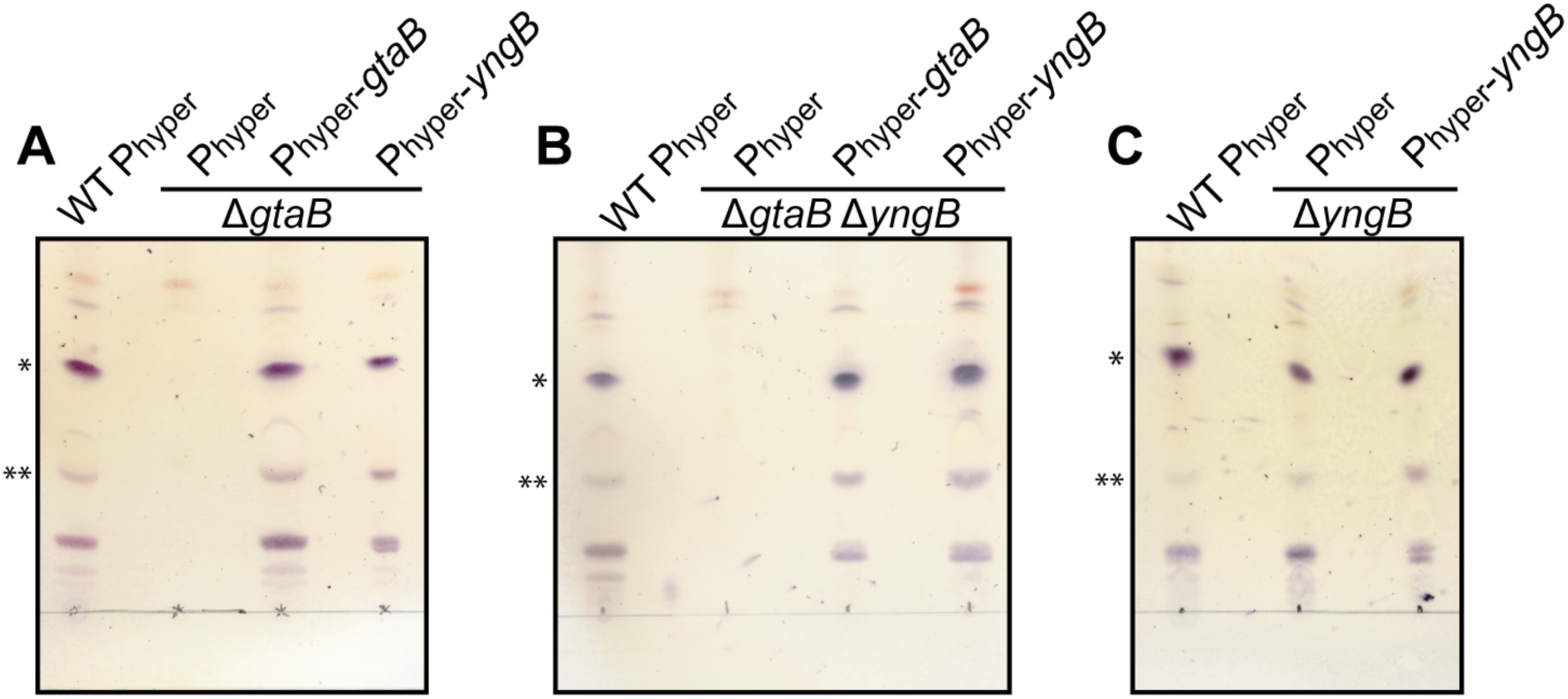
TLC analysis and detection of glycolipids produced by aerobically grown wild-type *B. subtilis*, mutant and complementation strains. Total membrane lipids were isolated from different *B. subtilis* strains following overnight growth under aerobic conditions in LB medium supplemented with IPTG and glucose. Lipids were separated by TLC and glycolipids visualized by spraying the plates with α-naphthol and 95% sulfuric acid and heating. Strains used were *A*, wild-type *B. subtilis* (WT P_hyper_), the *gtaB* mutant Δ*gtaB* P_hyper_, and complementation strains Δ*gtaB* P_hyper_-*gtaB* and Δ*gtaB* P_hyper_-*yngB*; *B*, wild-type *B. subtilis* (WT P_hyper_), the *gtaB/yngB* double mutant Δ*gtaB*Δ*yngB* P_hyper_ and complementation strains Δ*gtaB*Δ*yngB* P_hyper_-*gtaB* and Δ*gtaB*Δ*yngB* P_hyper_-*yngB*; and *C*, wild-type *B. subtilis* (WT P_hyper_), the *yngB* mutant Δ*yngB* P_hyper_, and complementation strain Δ*yngB* P_hyper_-*yngB*. Three independent experiments were performed, and a representative TLC plate image is shown. (*) marks likely Glc_2_-DAG and (**) likely Glc_3_-DAG glycolipid bands as assessed by MALDI-TOF mass spectrometry.

### YngB is produced under anaerobic growth condition, leading to decoration of WTA with glucose residues and glycolipid production

In previous work, it has been shown that expression of genes in the *yngABC* operon is upregulated by the transcription activator YclJ (35). Expression of *yclJ* itself is under control of the transcription factor ResD, which is produced under oxygen-limitation conditions (42–44). It is therefore possible that YngB is expressed from its native promoter under anaerobic growth conditions through a pathway involving ResD and YclJ. To determine if YngB is expressed from its native promoter under anaerobic growth condition and contributes to the decoration of WTA with glucose molecules and glycolipid production, the wild-type *B. subtilis* strain 168 and the isogenic Δ*gtaB* and Δ*yngB* single and Δ*gtaB*Δ*yngB* double mutant strains were grown in an anaerobic chamber under fermentative growth condition. The presence of glucose on WTA and the production of glycolipids was assessed by fluorescence microscopy and TLC analysis as described above. Clear fluorescence signals were observed for the wild-type, Δ*gtaB* mutant and Δ*yngB* mutant strains under anaerobic fermentative growth conditions, indicating that WTA is also decorated with glucose residues during anaerobic growth (Fig.6A). Only cells of the Δ*gtaB*Δ*yngB* double mutant were no longer stained (Fig.6A). These data suggest that under these growth conditions YngB is produced from its native promoter and functions as a second UGPase enzyme next to GtaB. Consistent with the fluorescence microscopy data, glycolipids could be detected by thin layer chromatography in wild-type, Δ*gtaB* and Δ*yngB* single mutant strains, but not in the Δ*gtaB*Δ*yngB* double mutant following growth under anaerobic, fermentative growth condition (Fig. 6B). Some differences in the glycolipid profiles were observed for lipid samples isolated from the wild-type following growth under aerobic or anaerobic conditions (Fig. 6B). A number of the slower migrating glycolipids were absent in samples derived from the anaerobically grown cultures, however bands likely corresponding to Glc_2_-DAG and Glc_3_-DAG were present in both samples (Fig. 6B). Furthermore, a reduced glycolipid signal was observed for the Δ*gtaB* mutant compared to the wild-type and Δ*yngB* mutant strain (Fig. 6B). This is consistent with the microscopy results and the observation that only some but not all of the Δ*gtaB* mutant cells seemed to contain glucose decorations on their WTA. Taken together, the data show that GtaB is the main UGPase enzyme producing UDP-glucose under both aerobic and anaerobic fermentative growth conditions. Furthermore, our data show that YngB is a functional UGPase that augments UDP-glucose production under oxygen limiting conditions. Under these conditions, it functions together with GtaB to produce UDP-glucose, for the production of glycolipids and the decoration of WTA with glucose residues and as discussed below potentially also other cell wall structures.

**Figure 6.**
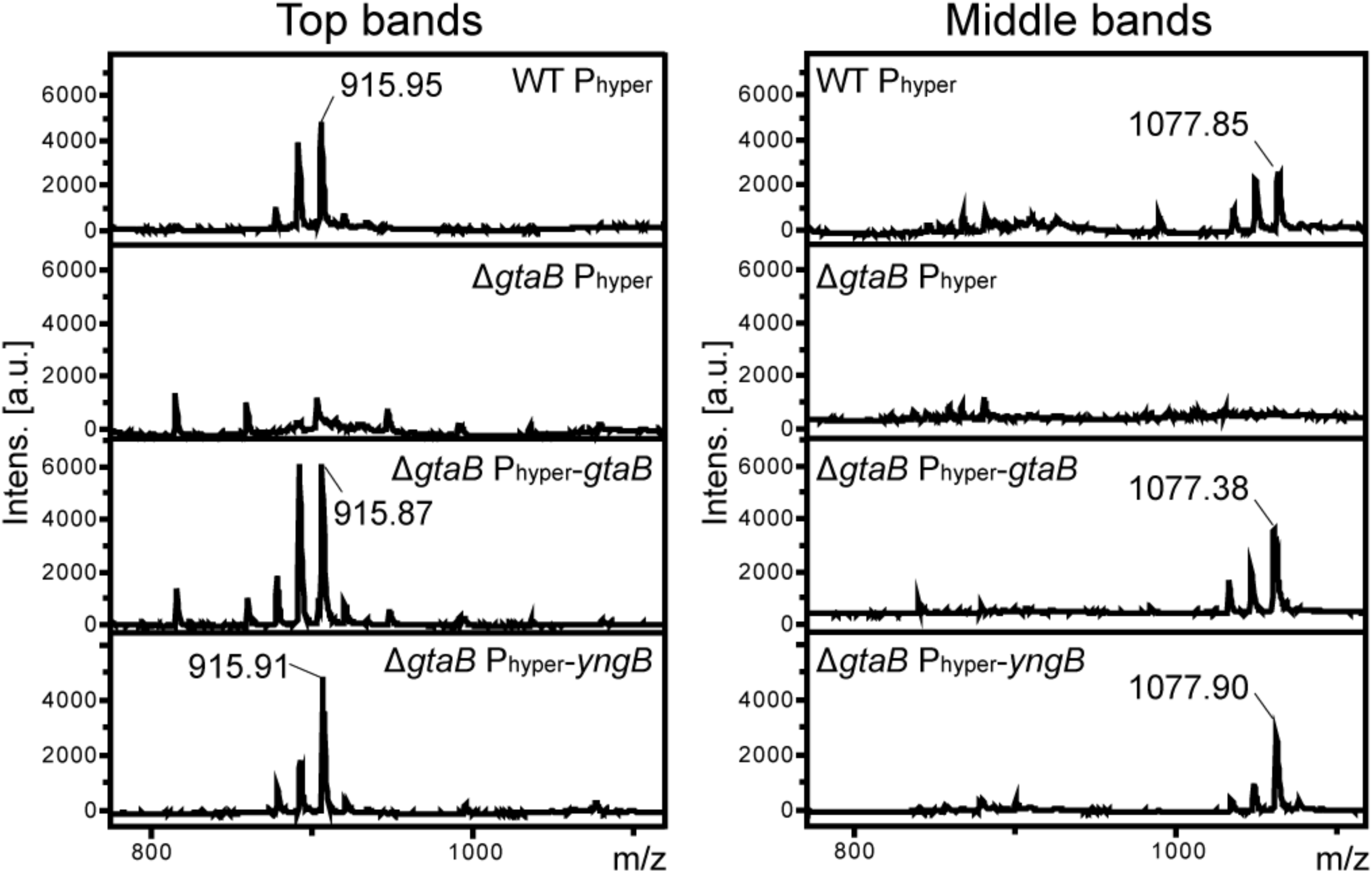
MALDI-TOF analysis of glycolipids isolated from wild-type *B. subtilis*, mutant and complementation strains. The glycolipid bands marked with * (top band) and ** (middle band) in Figure 5 were extracted from TLC plates for samples derived from wild-type *B. subtilis* (WT P_hyper_), the *gtaB* mutant Δ*gtaB* P_hyper_, and complementation strains Δ*gtaB* P_hyper_-*gtaB* and Δ*gtaB* P_hyper_-*yngB* and analyzed by MALDI-TOF M/S in the positive ion mode. Lipids were analyzed from 3 independent experiments and the spectra for one experiment is shown. The observed and expected masses of sodium adducts of specific glycolipids are summarized in Table 2. The data indicate that the top bands (*) corresponds to Glc_2_-DAG and the middle bands (**) to Glc_3_-DAG.

## Discussion

In bacteria, nucleotide-activated sugars are key sugar donors for glycosylation processes (52,55). In the Gram-positive, spore-forming bacterium *B. subtilis*, one of the best characterized nucleotide-activated sugar-synthesizing enzymes is GtaB. Up to now all UDP-glucose produced in *B. subtilis* has been attributed to the activity of GtaB, despite the presence of two orthologous proteins, YngB and YtdA. Here we show that, based on its crystal structure and *in vitro* and *in vivo* biochemical activity, the *B. subtilis* YngB protein is a functional UGPase (Figs 1 and 2). The necessary glucose-1-phosphate and UTP substrate-binding residues could be identified in the YngB structure (Fig. 1) and it is likely that YngB synthesizes UDP-glucose via the same catalytic mechanism proposed for other members of this family (47,48,52).

The main reason why all UDP-glucose production in *B. subtilis* has been attributed to GtaB is likely due to the specific growth conditions or developmental stages in which YngB and YtdA are produced. We present here experimental evidence that YngB is produced and can synthesize UDP-glucose when bacteria are grown under anaerobic conditions (Fig. 7). However, even under these conditions, GtaB appears to remain the main UGPase, as YngB activity could only be revealed in a *gtaB* mutant strain (Fig. 7). *ytdA* is under the control of the sporulation-specific transcription factor sigma K and hence only expressed during late stages of the sporulation process (56). It has been suggested that YtdA is a UGPase involved in the production of the polysaccharide layer on spores, however, no clear phenotype could be identified for a *ytdA* mutant strain (36). Therefore, similar as done here, to reveal a function of YtdA as UGPase, its activity and contribution to the production of the polysaccharide layer on spores might need to be assessed in a *gtaB* mutant strain.

**Figure 7.**
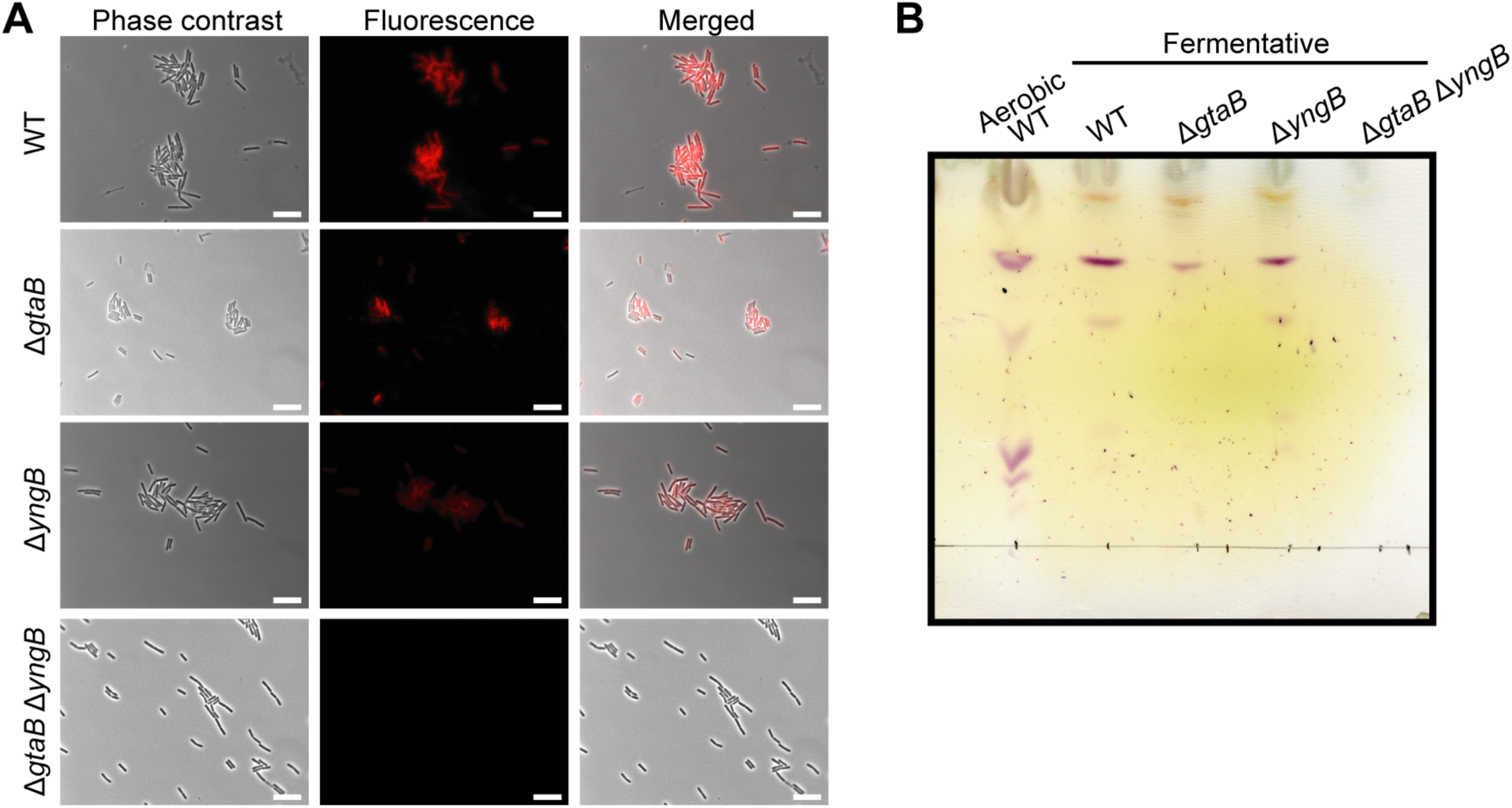
Detection of glucose modification on WTA and glycolipid analysis of wild-type and mutant *B. subtilis* strains following growth under anaerobic conditions. Wild-type *B. subtilis* (WT P_hyper_) and the *gtaB* and *yngB* single and double mutant strains Δ*gtaB* P_hyper_, Δ*yngB* P_hyper_, and Δ*gtaB*Δ*yngB* P_hyper_ were grown in an anaerobic chamber under fermentative growth condition on 2 x YT plates. *A*, bacteria were prepared for microscopy analysis and stained with the fluorescently labelled Alexa Fluor^™^ 594 Concanavalin A lectin to detect glucose modifications on WTA as described in the experimental procedure section. The experiment was performed 3 times and representative phase contrast, fluorescence and merged images are shown for each strain. Scale bars represent 10 µm. *B*, total membrane lipids were isolated from the different *B. subtilis* strains following growth in an anaerobic chamber, separated by TLC and glycolipids visualized by spraying the plates with α-naphthol and 95% sulfuric acid and heating. As control, a lipid sample isolated from the wild-type strain grown aerobically was run alongside the other samples. The experiment was performed 3 times and a representative TLC plate is shown.

Under aerobic growth condition, an aberrant morphology was observed for the Δ*gtaB* mutant; in contrast to rod-shaped wildtype bacteria, the mutant cells were curved (Fig. 3), consistent with previous observations (31,53,57). It is thought that the aberrant morphology is due to the lack of glycolipids, rather than the absence of glucose residues on WTA. This is based on the observation that an *ugtP* (*ypfP*) mutant unable to produce glycolipids but still containing glucose residues on WTA has an aberrant morphology, while a *tagE* mutant lacking glucose decorations on WTA but producing glycolipids does not display morphological defects (31,53,57). Consistent with these findings, the Δ*gtaB*Δ*yngB* double mutant strain, which is also unable to produce glycolipids, showed similar morphological defects under aerobic growth conditions (Figs 3 and 5). Interestingly, no aberrant morphology was observed for Δ*gtaB* or Δ*gtaB*Δ*yngB* mutant bacteria when grown under anaerobic growth conditions and both strains produced short, rod-shaped cells (Fig. 7). Under these growth conditions, the *gtaB* mutant is able to produce glycolipids, be it at reduced levels, while the Δ*gtaB*Δ*yngB* double mutant is unable to synthesize glycolipids (Fig. 7). These data not only show that YngB contributes to UDP-glucose production under anaerobic growth conditions, but they also indicate that the production of glycolipids is not essential for cells to maintain their normal rod-shape under anaerobic growth conditions.

The finding that YngB contributes to the production of UDP-glucose under anaerobic growth conditions is consistent with reports on its expression control. Transcription of the *yngABC* operon has been reported to be activated by YclJ, a transcription factor that is part of the YclJK two-component system, whose expression itself is upregulated during oxygen limitation by the ResDE two-component system (35,42,44). Previous studies have revealed the genes and operons regulated by YclJ, which include besides the *yngABC* operon, the two genes of the *ykcBC* operon (35,42) (Fig. 8A). Although we show here that the UDP-glucose produced by YngB can be utilized for the glucosylation of WTA and glycolipid production under anaerobic growth, given the predicted function of the proteins encoded by the *yngABC* and *ykcBC* operons, we speculate that the UDP-glucose produced by YngB could potentially be utilized for the glucosylation of other extracellular cell-wall components (Fig. 8). As outlined in detail below, this hypothesis is based on the observed similarities of the enzymes encodes in the *yngABC* and *ykcBC* operons to a multi-component transmembrane glycosylation system required for the extracellular glycosylation of LTA. In *B. subtilis*, WTA is glycosylated intracellularly by TagE, which adds the sugar residue directly onto the WTA backbone using a nucleotide-activated sugar as precursor, whereas LTA is glycosylated extracellularly using a multi-component transmembrane glycosylation system (21). Glycosylation of LTA starts when a nucleotide-activated sugar, in the case of *B. subtilis* UDP-GlcNAc, is linked by the glycosyltransferase CsbB to the C_55_-P lipid carrier in the cytosol of the cell (Fig. 8B). The C_55_-P-sugar intermediate is subsequently flipped across the membrane, likely by the GtrA-type membrane protein GtcA, and the sugar is finally added onto the LTA polymer on the outside of the cell by the multi-membrane spanning GT-C-type glycosyltransferase YfhO (Fig. 8B) (19–21). Proteins with similarity to CsbB, GtcA and YfhO are encoded in the YclJ-controlled *yngABC* and *ykcBC* operons. YkcC shows similarity to the glycosyltransferase CsbB and could therefore function to produce a C_55_-P-sugar intermediate (Fig. 8C). YngA is similar to GtcA, which is predicted to mediate the transport of C_55_-P-sugar intermediates across the membrane (Fig. 8C). Finally, YkcB is predicted to be a multi-membrane spanning GT-C-fold glycosyltransferase similar to YfhO and could therefore transfer the sugar residue from the exported C_55_-P-sugar intermediate onto an extracellular cell wall component (Fig. 8C). By analogy to the function of CsbB, GtcA and YfhO, we therefore speculate that YkcC, YngA and YkcB constitute a multi-component transmembrane glycosylation system that adds sugar decorations onto a cell wall component on the extracellular side of the membrane (Fig. 8C). We have shown here that YngB is a functional UGPase, hence we speculate that UDP-glucose is the likely substrate for the proposed YkcC-YngA-YkcB multi-component transmembrane glycosylation system (Fig. 8C). While the glycosylation target in the bacterial cell envelope is currently unknown, it will be interesting to investigate this further in future studies. As the transcription of the genes in the *yngABC* and *ykcBC* operons is predicted to be activated during anaerobic growth conditions, the YkcC-YngA-YkcB system likely functions only during specific growth conditions and therefore may mediate glycosylations that form a specific adaptation to these growth conditions. Furthermore, while UDP-glucose produced by YngB is utilized by TagE and UgtP, it will also be interesting to determine if, through specific protein/protein interactions, the UDP-glucose produced by YngB can be more efficiently fed towards the YkcC-YngA-YkcB multi-component transmembrane glycosylation system, as compared to TagE and UgtP, enzymes which are supplied with UDP-glucose by GtaB. Such specific protein/protein interactions that allow for a more efficient transfer of the soluble UDP-glucose precursor to specific enzymes, could be a reason why *B. subtilis* has evolved to produce not just one but multiple UGPases.

**Figure 8.**
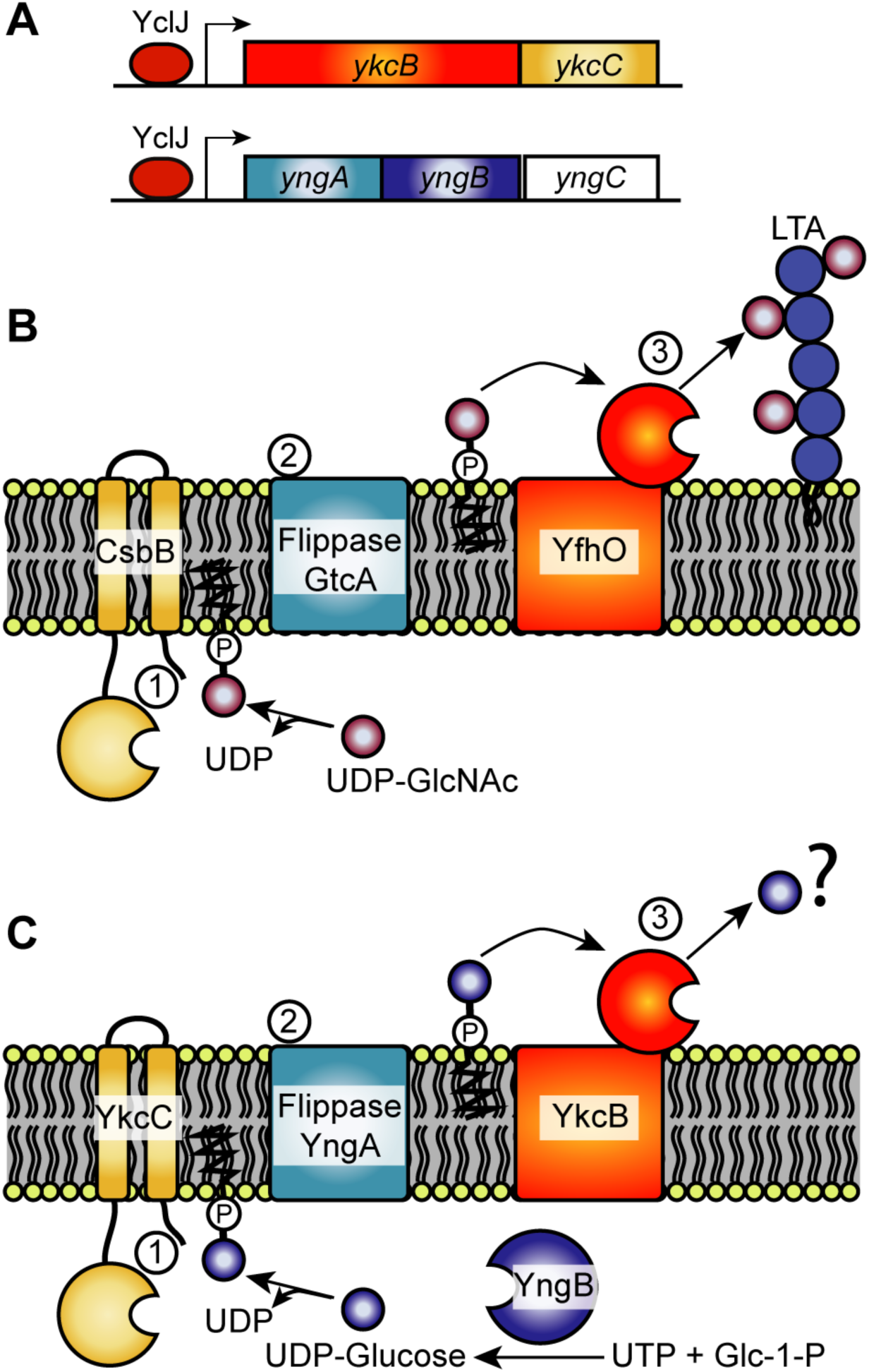
Proposed glycosylation pathway involving the UGPase YngB and the YkcC-YngA-YkcB multi-component transmembrane glycosylation system. *A*, the transcription factor YclJ activates the transcription of genes belonging to the *ykcBC* and *yngABC* operons. *B*, the *B. subtilis* LTA glycosylation model requires CsbB, GtcA and YfhO for the transfer of GlcNAc onto the LTA. The glycosyltransferase CsbB produces C_55_-P-GlcNAc, which is subsequently transported across the membrane by GtcA and the GlcNAc is finally attached to the LTA polymer on the outside of the cell by the multimembrane spanning GT-C-fold glycosyltransferase YfhO. *C*, proposed glycosylation pathway leading under anaerobic growth conditions to the transfer of glucose onto an uncharacterized target in the cell envelope. As shown in this study, YngB is a functional UGPase that can produce UDP-glucose and we hypothesize, that the predicted glycosyltransferase YkcC produces C_55_-P-glucose, which is transported across the membrane by YngA. The glucose residue is then transferred by the predicted multimembrane spanning GT-C-fold glycosyltransferase YkcB to an unknown target within the bacterial cell envelope. Panel B was adapted from model figure presented in Rismondo *et al*., 2018 (19) and Rismondo *et al*., 2020 (20).

In conclusion, we provide the first experimental evidence that the *B. subtilis* YngB protein is a functional UGPase that is produced under anaerobic growth conditions. The UDP-glucose synthesized by YngB is utilized for the glycosylation of WTA as well as glycolipid production. However, we speculate that the UDP-glucose produced by YngB might also enter other glycosylation pathways, leading to the decorating of other cell envelope components with glucose residues under anaerobic or other growth conditions, under which genes forming part of the YclJ regulon are activated.

## Experimental procedures

### Bacterial strains and growth conditions

All strains used in this study are listed in Table S2. *Escherichia coli* and *B. subtilis* strains were grown in Luria-Bertani (LB) medium at 37°C unless otherwise stated. LB medium for *B. subtilis* was supplemented with 0.2 % D-glucose for aerobic growth condition. For anaerobic fermentative growth of *B. subtilis*, a single colony was first inoculated in LB medium and grown at 37°C aerobically until reaching an OD_600_ of 1.0. Then 100 µl of the bacteria culture was spread on 2 × yeast extract tryptone (YT) agar plates (16g/L tryptone, 10g/L yeast extract, 5g/L NaCl, 1% glucose, 20 mM K_3_PO_4_ pH 7.0, 15g/L Bacto agar). The plates were incubated for 18 hr at 37°C in an anaerobic cabinet (Don Whitley Scientific) with an atmosphere of 10% CO_2_, 10% H_2_, and 80% N_2_. Bacterial cultures were supplemented with appropriate antibiotics at the following final concentrations: *E. coli* cultures, ampicillin (Amp) at 100 µg/ml and kanamycin (Kan) at 30 µg/ml; for *B. subtilis* cultures, Kan at 10 µg/ml, erythromycin (Erm) at 5 µg/ml, spectinomycin (Spec) at 100 µg/ml.

### Strain and plasmid construction

All primers used in this study are listed in Table S3. For the construction of plasmids for the expression and purification of the C-terminally His-tagged *B. subtilis* GtaB and YngB proteins, the *gtaB* (BSU_35670) and *yngB* (BSU_18180) genes were amplified by PCR from *B. subtilis* 168 genomic DNA using primer pairs ANG3161/ANG3162 and ANG3163/ANG3164, respectively. The PCR products were digested with NcoI and XhoI and ligated with plasmid pET28b cut with the same restriction enzymes. The resulting plasmids pET28b-*gtaB*-cHis and pET28b-*yngB*-cHis were recovered in *E. coli* strain XL1-Blue, yielding strains ANG5206 and ANG5207, respectively. Sequence of the inserts for pET28b plasmids was confirmed by sequencing using primers ANG111 and ANG112. For protein expression, the plasmids pET28b-*gtaB*-cHis and pET28b-*yngB*-cHis were introduced into *E. coli* strain BL21(DE3), yielding strains ANG5208 and ANG5209.

For the construction of *B. subtilis* expressing *gtaB* or *yngB* from the IPTG-inducible hyperspank promoter (P_hyper_), the *gtaB* (BSU_35670) and *yngB* (BSU_18180) genes were amplified by PCR using *B. subtilis* 168 genomic DNA as template and primer sets ANG3203/ANG3204 and ANG3205/ANG3206, respectively. The PCR products were digested with HindIII and NheI and ligated with plasmid pDR111 cut with the same restriction enzymes. The resulting plasmids pDR111-*gtaB* and pDR111-*yngB* were recovered in *E. coli* strain XL1-Blue, yielding strains XL1-Blue-pDR111-*gtaB* and XL1-Blue-pDR111-*yngB*. The sequences of the inserts in pDR111 plasmids were confirmed by sequencing using primers ANG1671 and ANG1672. Plasmids pDR111, pDR111-*gtaB*, and pDR111-*yngB* were linearized with ScaI and introduced into the wild-type *B. subtilis* strain 168 yielding 168 *amy::spec* P_hyper_ (ANG5675), 168 *amy::spec* P_hyper_*-gtaB* and 168 *amy::spec* P_hyper_*-yngB* respectively. Next, the chromosomal DNA of strain 168Δ*gtaB::kan* (ANG5277) was introduced to strains 168 *amy::spec* P_hyper_, 168 *amy::spec* P_hyper_*-gtaB* and 168 *amy::spec* P_hyper_*-yngB* yielding strains 168Δ*gtaB::kan amy::spec* P_hyper_ (ANG5676), 168Δ*gtaB::kan amy::spec* P_hyper_*-gtaB* (ANG5677) and 168Δ*gtaB::kan amy::spec* P_hyper_*-yngB* (ANG5678), respectively. The chromosomal DNA of strain 168Δ*yngB::kan* (ANG5263) was introduced into 168 *amy::spec* P_hyper_ and 168 *amy::spec* P_hyper_*-yngB* yielding strains 168Δ*yngB::kan amy::spec* P_hyper_ (ANG5679) and 168Δ*yngB::kan amy::spec* P_hyper_*-yngB* (ANG5680). The chromosomal DNA of strain 168Δ*yngB::erm* (ANG5659) was introduced into strain ANG5677, yielding strain 168Δ*gtaB::kan* Δ*yngB::erm amy::spec* P_hyper_*-gtaB*(ANG5682). The chromosomal DNA of strain 168Δ*gtaB::erm* (ANG5658) was introduced into strains ANG5679 and ANG5680, yielding strains 168Δ*gtaB::erm* Δ*yngB::kan amy::spec* P_hyper_ (ANG5681) and 168Δ*gtaB::erm* Δ*yngB::kan amy::spec* P_hyper_*-yngB* (ANG5683). The deletion of the *gtaB* and *yngB* genes were confirmed by PCR using primer sets ANG3197/ANG3198 and ANG3199/ANG3200, respectively. The integration of the *gtaB* gene at the *amyE* site was confirmed by PCR using primer sets ANG1663/ANG3204 and ANG1664/ANG3203. The integration of the *yngB* gene at the *amyE* site was confirmed by PCR using primer sets ANG1663/ANG3206 and ANG1664/ANG3205. The integration of plasmid pDR111 at the *amyE* site for strain 168 *amy::spec* P_hyper_ was confirmed by PCR using primers sets ANG1664/ANG1671 and ANG1663/ANG1672.

### Expression and purification of GtaB and YngB

*E. coli* strain BL21(DE3) pET28b-*gtaB*-cHis (ANG5208) was grown in LB medium at 30°C with shaking until reaching an OD_600_ of 0.6. Protein expression was induced by the addition of IPTG to a final concentration of 0.5 mM and the cultures were incubated overnight at 16 °C with agitation. Bacterial cells were harvested by centrifugation and washed once with cold 500mM NaCl, 50mM Tris pH 7.5 buffer and the bacterial pellets stored at −20°C for future use. For the protein purification, the bacterial cells were suspended in 20 ml cold buffer A (500mM NaCl, 50mM Tris pH 7.5, 5% glycerol, 10 mM imidazole) supplemented with cOmplete^™^ protease inhibitor cocktail (Roche), 100 ug/ml lysozyme, 10 ug/ml DNase, followed by passing the cell suspension twice through a French press cell at 1100 psi. For the purification of GtaB-cHis, the cell lysate was loaded onto a 5ml HisTrap column equilibrated with buffer A. The column was washed with 5 column volumes of buffer A followed by elution using a linear gradient of 10 column volumes from buffer A to buffer B (500mM NaCl, 50mM Tris pH 7.5, 5% glycerol, 500 mM imidazole). Elution fractions containing GtaB-cHis were pooled and thrombin was added to cleave off the C-terminal His-tag. The protein was dialyzed at room temperature against 1L buffer C (500mM NaCl, 50mM Tris pH 7.5, 5% glycerol) for 1 hr, followed by overnight dialysis against 1L of fresh buffer C. The protein solution was then loaded onto a Superdex 10/60 Hiload size exclusion column (GE Healthcare) equilibrated with buffer C. Following size-exclusion chromatography, the purified protein was concentrated using a PES 10-kDa cut off Pierce^™^ protein concentrator.

For the expression of selenomethionine substituted YngB-cHis, strain BL21(DE3) pET28b-*yngB*-cHis (ANG 5209) was grown in LB medium at 37° for 12 hr. Bacterial cells were then washed once with minimal medium (0.5 g/L NaCl, 1.0 g/L (NH_4_)_2_SO_4_, 7.5 g/L KH_2_PO_4_, 23.25 g/L K_2_HPO_4_, 0.246 g/L MgSO_4_, 7.2 g/L glucose, 100 mg/L lysine, 100 mg/L phenylalanine, 100 mg/L threonine, 50 mg/L isoleucine, 50 mg/L leucine, 50 mg/L valine, 0.1 mM CaCl_2_, 10 ml/L 100 × Kao and Michayluk vitamin solution) and grown overnight at 37°C in the minimal medium containing 42 mg/L methionine. The next day, bacterial cells were washed with the minimal medium and grown in the minimal medium containing 42 mg/L selenomethionine at 37°C until reaching an OD_600_ of 0.5 – 0.6. At this point 2.9 g/L additional glucose and IPTG to give a final concentration of 0.5 mM, were added. The bacterial culture was incubated at 16°C overnight with agitation. Cells were harvested, washed and stored as described above. For the production of native YngB-cHis used in the kinetic assay, methionine was added to the medium in place of selenomethionine. For the purification of SeMet YngB-cHis or native YngB-cHis, the cell lysate was loaded by gravity flow onto a column containing 1 ml Ni-NTA resin (Qiagen) equilibrated with buffer A. The column was washed with 30 ml of buffer A and 30 ml of buffer D (500mM NaCl, 50mM Tris pH 7.5, 5% glycerol, 50 mM imidazole). The proteins were eluted in 5 × 1 ml fractions using buffer B. The elution fractions were pooled and subjected to size-exclusion chromatography and the purified protein concentrated as described above.

### Protein crystallization, structural solution and analysis

SeMet YngB-cHis crystals were obtained by the sitting drop-method in 0.2M potassium citrate tribasic monohydrate, 0.05M lithium citrate tribasic tetrahydrate, 0.1M sodium phosphate monobasic monohydrate, 25% PEG6000, using a protein concentration of 6 mg/ml. SeMet YngB-cHis crystals were cryo-protected with 30% ethylene glycol and flash frozen in liquid nitrogen. Datasets were collected at the I03 Beamline at the Diamond Light Source (Harwell Campus, Didcot, UK). Data indexing, integration, scaling, and merging was done using the xia2 3dii pipeline (58). The selenium sites and initial phases were solved using CRANK2 (59). Structure refinement was performed with Refmac (60) and model building with Coot (61). Data collection and refinement statistics are summarized in Table 1. The structure was validated through the Validation Pipeline wwPDB-VP and the geometry outliers assessed using Molprobity (62).The figures with the structure were generated using PyMol.

### Enzyme kinetic analysis

The steady state kinetics assays with purified GtaB and YngB-cHis proteins were performed using a previously described method with some modifications (52). Briefly, enzyme assays were performed in 96-well plates in 100 µl reaction volumes. The reactions contained 50 mM Tris pH 7.5, 500 mM NaCl, 5 % glycerol, 10 mg/ml MgCl_2_, 0.05U of *E. coli* pyrophosphatase (NEB), 100 nM GtaB_Bs_ or 100 nM YngB_Bs_-cHis. For measuring the K_m_ value for α-glucose-1-phosphate (G-1-P), the reactions contained 200 µM UTP and α-glucose-1-phosphate at a concentration of 200 µM, 150 µM, 100 µM, 50 µM, 12.5 µM, 6.3 µM, or 3.1 µM. For measuring the K_m_ value of UTP, the reactions contained 200 µM α-glucose-1-phosphate and UTP at a concentration of 200 µM, 150 µM, 100 µM, 50 µM, 12.5 µM, 6.3 µM, or 3.1 µM. Reactions were performed in triplicate and reactions without GtaB_Bs_ and YngB_Bs_-cHis were used as negative controls. Reactions were incubated at room temperature for 1 min for GtaB_Bs_ and 4 min for YngB_Bs_-cHis so that less than 20% of the substrates were converted in the reactions. The reactions were terminated by adding 100 µl of Biomol^®^ Green (Enzo^®^ Life Sciences) and after a 20 min incubation at room temperature, the absorbance was measured at 620 nm. The number of UDP-glucose formed per molecule of GtaB or YngB per second is plotted versus substrate concentration, followed by Michaelis-Menten non-linear fitting using Prism.

### Detection of glucose residues on WTA using fluorescently labelled lectin Concanavalin A

Glucose modifications on WTA were detected by fluorescence microscopy using a previously described method with minor modifications (34). Single colonies of the different *B. subtilis* strains were used to inoculate 5 ml LB medium and the cultures grown overnight at 37°C. The overnight cultures were back diluted 1:100 into 25 ml of fresh LB medium supplemented with 1 mM IPTG and the cultures grown at 37°C until reaching mid-log growth phase (OD_600_ between 0.4 and 0.6). Cells equivalent to 100 µl of a culture with an OD_600_ of 0.5 were pelleted by centrifugation at 17,000 × g for 1 min and washed once with PBS pH 7.4. Bacterial cells were subsequently suspended in 80 µl PBS and mixed with 20 µl of 1 mg/ml Concanavalin A AlexaFluor^™^ 594 conjugate dissolved in 0.1 M sodium bicarbonate pH 8.3, followed by incubation at room temperature for 30 min in the dark. The bacterial cells were subsequently washed three times with 100 µl PBS and 1min centrifugation steps to pellet cells and finally suspended in 100 µl PBS. Microscopic analysis was performed as previously described (20). Briefly, samples were spotted on microscope slides coated with a thin agarose film (1.2% agarose in distilled water). Phase contrast and fluorescence images were taken using a 100x objective and a Zeiss Axio Imager.A1 microscope coupled to the AxioCam MRm and processed using the Zen 2012 (blue edition) software. The Zeiss filter set 00 was used for the detection of fluorescence signals. For the microscopy experiment using bacteria grown under anaerobic fermentative growth conditions, colonies obtained on agar plates incubated for 18 h at 37°C in an anaerobic cabinet were suspended in 100 µl PBS to an OD_600_ of 0.5. The staining and microscopic analysis was performed as described above. Representative data from three independent experiments are shown.

### Isolation of WTA and its analysis by NMR and UPLC-MS

*B. subtilis* strains were grown in 2 L LB medium supplemented with 1mM IPTG at 37 °C. Once the cultures reached an OD_600_ of 0.6, the cells were harvested by centrifugation and WTA isolated using a previously described method (19). The NMR analysis of WTA was performed as previously described (19,20). Briefly, 2 mg of WTA from each strain was suspended and lyophilized twice in 500 μl D_2_O of 99.96% purity. Lyophilized WTA at the final step was suspended in 500 μl D_2_O of 99.96% purity and NMR spectra were recorded on a 600-MHz Bruker Advance III spectrometer equipped with a TCI cryoprobe. NMR spectra were recorded at 303 K with a total recycling time of 5 s and a ^1^H flip angle of approximately 30°. Two independent experiments were performed, and very similar spectra were obtained and one spectrum for each strain is shown. For the UPLC-MS analysis of the purified WTA, the method used was adapted from previously described protocols (20,63). Briefly, 2 mg of the purified WTA was lyophilized in deionized distilled H_2_O. The lyophilized WTA was then depolymerized into monomeric units by hydrolysis of the phosphodiester bonds using 48 % hydrofluoric acid for 20 hr at 0°C. The depolymerized WTA material was subjected to UPLC-MS analysis as described previously (63). All data were collected and processed using the MassLynx software, version 4.1 (Waters Corp., USA).

### Isolation of membrane lipids and TLC analysis

For the isolation of total membrane lipids, the different *B. subtilis* strains were grown overnight at 37°C in 100 ml LB medium supplemented with 1 mM IPTG. Cells were harvested by centrifugation, and total membrane lipids isolated as described previously (64). TLC analysis and detection of glycolipids was performed as described (64). Briefly, isolated lipids were suspended in chloroform and 0.5 mg were spotted on Å60 silica gel plates (Macherey-Nagel). Lipids were separated using a developing solvent of chloroform: methanol: H_2_O (65:25:4). Plates were sprayed with 0.5% α-naphthol in 50% methanol and then with 95% sulfuric acid. Glycolipids were visualized as purple bands by a final heating step. For the glycolipid analysis of bacteria grown under anaerobic fermentative conditions, colonies obtained on agar plates following incubated at 37°C in an anaerobic cabinet were scraped off the plates and suspended in 0.1M sodium citrate pH 4.7 for subsequent membrane lipid isolation and TLC analysis as described above. Representative data from three independent experiments are shown.

### MALDI-TOF MS and MALDI-TOF MS/MS

#### analysis of glycolipids

MALDI-TOF analysis of glycolipids was performed using a previously described method with some minor modifications (64). A total of 5 × 0.5 mg of lipids were spotted on silica plates and separated by TLC as described above. The silica matrix with the lipids from appropriate areas of the TLC plates were scraped into glass tubes and extracted overnight at room temperature with 6 ml of a 1:1 methanol: chloroform mix for each sample. Next day, the silica matrix was removed by filtering the solutions through classic Sep-pak silica cartridges (Waters) pre-equilibrated with 6 ml methanol followed by 6 ml chloroform. The filtered samples with the extracted lipids were dried under a stream of nitrogen. Dried lipids were suspended in 50 µl chloroform and aliquots mixed 1:1 with matrix. The matrix consisted of a 9:1 mixture of 2,5-dihydroxybenzoic acid and 2-hydroxy-5-methoxybenzoic acid (super-DHB, Sigma-Aldrich) at a final concentration of 10 mg/ml dissolved in chloroform: methanol at a ratio of 9:1. One µl sample was spotted onto disposable MSP 96 polished steel plate. As calibration standard, the peptide calibration standard II (Bruker) in 0.1% TFA was mixed 1:1 with IVD Matrix α-Cyano-4-hydroxycinnamic acid. The samples were analyzed on a MALDI Biotyper Sirius system (Bruker Daltonik, Germany). The mass profiles were acquired using the FlexControl 3.4 software (Bruker Daltonik, Germany) with mass spectra scanned in the m/z range of 600 to 2,000. Spectra were recorded in the reflector positive ion mode (laser intensity 95%, ion source 1 = 10.00 kV, ion source 2 = 8.98 kV, lens = 3.00 kV, detector voltage = 2652 V, pulsed ion extraction = 150 ns). Each spectrum corresponded to an ion accumulation of 5,000 laser shots randomly distributed on the spot. Representative spectra from two independent experiments are shown. The obtained spectra were processed with default parameters using the FlexAnalysis v.3.4 software (Bruker Daltonik, Germany). For the MALDI-TOF MS/MS analysis, MS/MS fragmentation profiles were acquired on a 4800 Proteomics Analyzer (with TOF-TOF Optics, Applied Biosystems, plate: 384 Opti-TOF 123 mm × 84 mm AB Sciex NC0318050, 1016629) using the reflectron mode. Samples were analyzed operating at 20 kV in the positive ion mode. MS/MS mass spectrometry data were analyzed using the Data Explorer software version 4.9 from Applied Biosystems.

## Data availability

The atomic coordinates and structure factors have been deposited in the Protein Data Bank under accession code 7B1R. https://doi.org/10.2210/pdb7B1R/pdb

## Author contribution statement

**Chih-Hung Wu:** Conceptualization, Investigation, Data analysis, Visualization, Writing – original draft preparation. **Jeanine Rismondo:** Investigation, Supervision, Data analysis, Writing – review & editing. **Rhodri M. L. Morgan:** Data analysis, Supervision, Writing – review & editing. **Yang Shen:** Investigation, Data analysis, Writing – review & editing. **Martin J. Loessner:** Data analysis, Writing – review & editing. **Gerald Larrouy-Maumus:** Investigation, Data analysis, Writing – review & editing **Paul S. Freemont:** Conceptualization, Funding acquisition, Supervision, Data analysis, Writing – review & editing. **Angelika Gründling:** Conceptualization, Funding acquisition, Data analysis, Supervision, Writing – original draft preparation.

## Acknowledgments

We thank Dr. Samy Bulous from the Institute of Food, Nutrition and Health, ETH Zürich, for the technical assistance with the UPLC-MS analysis.

## Funding and additional information

This work was funded by the MRC grant MR/P011071/1 to AG and PSF and the German research foundation (DFG) grant RI 2920/1-1 to JR. The crystallization facility at Imperial College was funded by the BBSRC (BB/D524840/1) and the Wellcome Trust (202926/Z/16/Z). Datasets were collected at the I03 Beamline at the Diamond Light Source (Harwell Campus, Didcot, UK).

## References

1. Wagner, J. K., Marquis, K. A., and Rudner, D. Z. (2009) SirA enforces diploidy by inhibiting the replication initiator DnaA during spore formation in Bacillus subtilis. Mol Microbiol 73, 963–974

2. Vollmer, W., Blanot, D., and de Pedro, M. A. (2008) Peptidoglycan structure and architecture. FEMS Microbiol Rev 32, 149–167

3. Cress, B. F., Englaender, J. A., He, W., Kasper, D., Linhardt, R. J., and Koffas, M. A. (2014) Masquerading microbial pathogens: capsular polysaccharides mimic host-tissue molecules. FEMS Microbiol Rev 38, 660–697

4. Whitfield, C., and Trent, M. S. (2014) Biosynthesis and export of bacterial lipopolysaccharides. Annu Rev Biochem 83, 99–128

5. Weidenmaier, C., and Peschel, A. (2008) Teichoic acids and related cell-wall glycopolymers in Gram-positive physiology and host interactions. Nat Rev Microbiol 6, 276–287

6. Missiakas, D., and Schneewind, O. (2017) Assembly and Function of the Bacillus anthracis S-Layer. Annu Rev Microbiol 71, 79–98

7. Brown, S., Santa Maria, J. P., Jr., and Walker, S. (2013) Wall teichoic acids of gram-positive bacteria. Annu Rev Microbiol 67, 313–336

8. Percy, M. G., and Gründling, A. (2014) Lipoteichoic acid synthesis and function in gram-positive bacteria. Annu Rev Microbiol 68, 81–100

9. Iwasaki, H., Shimada, A., Yokoyama, K., and Ito, E. (1989) Structure and glycosylation of lipoteichoic acids in Bacillus strains. J Bacteriol 171, 424–429

10. Young, F. E. (1967) Requirement of glucosylated teichoic acid for adsorption of phage in Bacillus subtilis 168. Proc Natl Acad Sci U S A 58, 2377–2384

11. Mauel, C., Young, M., Monsutti-Grecescu, A., Marriott, S. A., and Karamata, D. (1994) Analysis of Bacillus subtilis tag gene expression using transcriptional fusions. Microbiology 140 (Pt 9), 2279–2288

12. Lahooti, M., and Harwood, C. R. (1999) Transcriptional analysis of the Bacillus subtilis teichuronic acid operon. Microbiology 145 (Pt 12), 3409–3417

13. Lairson, L. L., Henrissat, B., Davies, G. J., and Withers, S. G. (2008) Glycosyltransferases: structures, functions, and mechanisms. Annu Rev Biochem 77, 521–555

14. Berbis, M. A., Sanchez-Puelles, J. M., Canada, F. J., and Jimenez-Barbero, J. (2015) Structure and Function of Prokaryotic UDP-Glucose Pyrophosphorylase, A Drug Target Candidate. Curr Med Chem 22, 1687–1697

15. Barreteau, H., Kovac, A., Boniface, A., Sova, M., Gobec, S., and Blanot, D. (2008) Cytoplasmic steps of peptidoglycan biosynthesis. FEMS Microbiol Rev 32, 168–207

16. Mengin-Lecreulx, D., and van Heijenoort, J. (1994) Copurification of glucosamine-1-phosphate acetyltransferase and N-acetylglucosamine-1-phosphate uridyltransferase activities of Escherichia coli: characterization of the glmU gene product as a bifunctional enzyme catalyzing two subsequent steps in the pathway for UDP-N-acetylglucosamine synthesis. J Bacteriol 176, 5788–5795

17. Mancuso, D. J., and Chiu, T. H. (1982) Biosynthesis of glucosyl monophosphoryl undecaprenol and its role in lipoteichoic acid biosynthesis. J Bacteriol 152, 616–625

18. Yokoyama, K., Araki, Y., and Ito, E. (1988) The function of galactosyl phosphorylpolyprenol in biosynthesis of lipoteichoic acid in Bacillus coagulans. Eur J Biochem 173, 453–458

19. Rismondo, J., Percy, M. G., and Gründling, A. (2018) Discovery of genes required for lipoteichoic acid glycosylation predicts two distinct mechanisms for wall teichoic acid glycosylation. J Biol Chem 293, 3293–3306

20. Rismondo, J., Haddad, T. F. M., Shen, Y., Loessner, M. J., and Gründling, A. (2020) GtcA is required for LTA glycosylation in Listeria monocytogenes serovar 1/2a and Bacillus subtilis. Cell Surf 6, 100038

21. Percy, M. G., Karinou, E., Webb, A. J., and Grundling, A. (2016) Identification of a Lipoteichoic Acid Glycosyltransferase Enzyme Reveals that GW-Domain-Containing Proteins Can Be Retained in the Cell Wall of Listeria monocytogenes in the Absence of Lipoteichoic Acid or Its Modifications. J Bacteriol 198, 2029–2042

22. Young, F. E., Smith, C., and Reilly, B. E. (1969) Chromosomal location of genes regulating resistance to bacteriophage in Bacillus subtilis. J Bacteriol 98, 1087–1097

23. Allison, S. E., D’Elia, M. A., Arar, S., Monteiro, M. A., and Brown, E. D. (2011) Studies of the genetics, function, and kinetic mechanism of TagE, the wall teichoic acid glycosyltransferase in Bacillus subtilis 168. J Biol Chem 286, 23708–23716

24. Genevaux, P., Bauda, P., DuBow, M. S., and Oudega, B. (1999) Identification of Tn10 insertions in the rfaG, rfaP, and galU genes involved in lipopolysaccharide core biosynthesis that affect Escherichia coli adhesion. Arch Microbiol 172, 1–8

25. Chang, H. Y., Lee, J. H., Deng, W. L., Fu, T. F., and Peng, H. L. (1996) Virulence and outer membrane properties of a galU mutant of Klebsiella pneumoniae CG43. Microb Pathog 20, 255–261

26. Bonofiglio, L., Garcia, E., and Mollerach, M. (2005) Biochemical characterization of the pneumococcal glucose 1-phosphate uridylyltransferase (GalU) essential for capsule biosynthesis. Curr Microbiol 51, 217–221

27. Cools, F., Torfs, E., Vanhoutte, B., de Macedo, M. B., Bonofiglio, L., Mollerach, M., Maes, L., Caljon, G., Delputte, P., Cappoen, D., and Cos, P. (2018) Streptococcus pneumoniae galU gene mutation has a direct effect on biofilm growth, adherence and phagocytosis in vitro and pathogenicity in vivo. Pathog Dis 76

28. Pooley, H. M., Paschoud, D., and Karamata, D. (1987) The gtaB marker in Bacillus subtilis 168 is associated with a deficiency in UDPglucose pyrophosphorylase. J Gen Microbiol 133, 3481–3493

29. Yasbin, R. E., Maino, V. C., and Young, F. E. (1976) Bacteriophage resistance in Bacillus subtilis 168, W23, and interstrain transformants. J Bacteriol 125, 1120–1126

30. Jorasch, P., Wolter, F. P., Zahringer, U., and Heinz, E. (1998) A UDP glucosyltransferase from Bacillus subtilis successively transfers up to four glucose residues to 1,2-diacylglycerol: expression of ypfP in Escherichia coli and structural analysis of its reaction products. Mol Microbiol 29, 419–430

31. Matsuoka, S., Seki, T., Matsumoto, K., and Hara, H. (2016) Suppression of abnormal morphology and extracytoplasmic function sigma activity in Bacillus subtilis ugtP mutant cells by expression of heterologous glucolipid synthases from Acholeplasma laidlawii. Biosci Biotechnol Biochem 80, 2325–2333

32. Matsuoka, S. (2018) Biological functions of glucolipids in Bacillus subtilis. Genes Genet Syst 92, 217–221

33. Luo, Y. (2016) Alanylated lipoteichoic acid primer in Bacillus subtilis. F1000Res 5, 155

34. Habusha, M., Tzipilevich, E., Fiyaksel, O., and Ben-Yehuda, S. (2019) A mutant bacteriophage evolved to infect resistant bacteria gained a broader host range. Mol Microbiol 111, 1463–1475

35. Ogura, M., Tsukahara, K., and Tanaka, T. (2010) Identification of the sequences recognized by the Bacillus subtilis response regulator YclJ. Arch Microbiol 192, 569–580

36. Shuster, B., Khemmani, M., Nakaya, Y., Holland, G., Iwamoto, K., Abe, K., Imamura, D., Maryn, N., Driks, A., Sato, T., and Eichenberger, P. (2019) Expansion of the Spore Surface Polysaccharide Layer in Bacillus subtilis by Deletion of Genes Encoding Glycosyltransferases and Glucose Modification Enzymes. J Bacteriol 201

37. Eiamphungporn, W., and Helmann, J. D. (2008) The Bacillus subtilis sigma(M) regulon and its contribution to cell envelope stress responses. Mol Microbiol 67, 830–848

38. Panta, P. R., Kumar, S., Stafford, C. F., Billiot, C. E., Douglass, M. V., Herrera, C. M., Trent, M. S., and Doerrler, W. T. (2019) A DedA Family Membrane Protein Is Required for Burkholderia thailandensis Colistin Resistance. Front Microbiol 10, 2532

39. Thompkins, K., Chattopadhyay, B., Xiao, Y., Henk, M. C., and Doerrler, W. T. (2008) Temperature sensitivity and cell division defects in an Escherichia coli strain with mutations in yghB and yqjA, encoding related and conserved inner membrane proteins. J Bacteriol 190, 4489–4500

40. Doerrler, W. T., Sikdar, R., Kumar, S., and Boughner, L. A. (2013) New functions for the ancient DedA membrane protein family. J Bacteriol 195, 3–11

41. Fabret, C., Feher, V. A., and Hoch, J. A. (1999) Two-component signal transduction in Bacillus subtilis: how one organism sees its world. J Bacteriol 181, 1975–1983

42. Kobayashi, K., Ogura, M., Yamaguchi, H., Yoshida, K., Ogasawara, N., Tanaka, T., and Fujita, Y. (2001) Comprehensive DNA microarray analysis of Bacillus subtilis two-component regulatory systems. J Bacteriol 183, 7365–7370

43. Ye, R. W., Tao, W., Bedzyk, L., Young, T., Chen, M., and Li, L. (2000) Global gene expression profiles of Bacillus subtilis grown under anaerobic conditions. J Bacteriol 182, 4458–4465

44. Härtig, E., Geng, H., Hartmann, A., Hubacek, A., Munch, R., Ye, R. W., Jahn, D., and Nakano, M. M. (2004) Bacillus subtilis ResD induces expression of the potential regulatory genes yclJK upon oxygen limitation. J Bacteriol 186, 6477–6484

45. Sun, G., Sharkova, E., Chesnut, R., Birkey, S., Duggan, M. F., Sorokin, A., Pujic, P., Ehrlich, S. D., and Hulett, F. M. (1996) Regulators of aerobic and anaerobic respiration in Bacillus subtilis. J Bacteriol 178, 1374–1385

46. Nakano, M. M., Zuber, P., Glaser, P., Danchin, A., and Hulett, F. M. (1996) Two-component regulatory proteins ResD-ResE are required for transcriptional activation of fnr upon oxygen limitation in Bacillus subtilis. J Bacteriol 178, 3796–3802

47. Kim, H., Choi, J., Kim, T., Lokanath, N. K., Ha, S. C., Suh, S. W., Hwang, H. Y., and Kim, K. K. (2010) Structural basis for the reaction mechanism of UDP-glucose pyrophosphorylase. Mol Cells 29, 397–405

48. Thoden, J. B., and Holden, H. M. (2007) Active site geometry of glucose-1-phosphate uridylyltransferase. Protein Sci 16, 1379–1388

49. Hanukoglu, I. (2015) Proteopedia: Rossmann fold: A beta-alpha-beta fold at dinucleotide binding sites. Biochem Mol Biol Educ 43, 206–209

50. Thoden, J. B., and Holden, H. M. (2007) The molecular architecture of glucose-1-phosphate uridylyltransferase. Protein Sci 16, 432–440

51. Aragao, D., Fialho, A. M., Marques, A. R., Mitchell, E. P., Sa-Correia, I., and Frazao, C. (2007) The complex of Sphingomonas elodea ATCC 31461 glucose-1-phosphate uridylyltransferase with glucose-1-phosphate reveals a novel quaternary structure, unique among nucleoside diphosphate-sugar pyrophosphorylase members. J Bacteriol 189, 4520–4528

52. Benini, S., Toccafondi, M., Rejzek, M., Musiani, F., Wagstaff, B. A., Wuerges, J., Cianci, M., and Field, R. A. (2017) Glucose-1-phosphate uridylyltransferase from Erwinia amylovora: Activity, structure and substrate specificity. Biochim Biophys Acta Proteins Proteom 1865, 1348–1357

53. Lazarevic, V., Soldo, B., Medico, N., Pooley, H., Bron, S., and Karamata, D. (2005) Bacillus subtilis alpha-phosphoglucomutase is required for normal cell morphology and biofilm formation. Appl Environ Microbiol 71, 39–45

54. Matsuoka, S., Chiba, M., Tanimura, Y., Hashimoto, M., Hara, H., and Matsumoto, K. (2011) Abnormal morphology of Bacillus subtilis ugtP mutant cells lacking glucolipids. Genes Genet Syst 86, 295–304

55. Horecker, B. L. (1966) The biosynthesis of bacterial polysaccharides. Annu Rev Microbiol 20, 253–290

56. Arrieta-Ortiz, M. L., Hafemeister, C., Bate, A. R., Chu, T., Greenfield, A., Shuster, B., Barry, S. N., Gallitto, M., Liu, B., Kacmarczyk, T., Santoriello, F., Chen, J., Rodrigues, C. D., Sato, T., Rudner, D. Z., Driks, A., Bonneau, R., and Eichenberger, P. (2015) An experimentally supported model of the Bacillus subtilis global transcriptional regulatory network. Mol Syst Biol 11, 839

57. Price, K. D., Roels, S., and Losick, R. (1997) A Bacillus subtilis gene encoding a protein similar to nucleotide sugar transferases influences cell shape and viability. J Bacteriol 179, 4959–4961

58. Winter, G., Lobley, C. M., and Prince, S. M. (2013) Decision making in xia2. Acta Crystallogr D Biol Crystallogr 69, 1260–1273

59. Skubak, P., and Pannu, N. S. (2013) Automatic protein structure solution from weak X-ray data. Nat Commun 4, 2777

60. Kovalevskiy, O., Nicholls, R. A., Long, F., Carlon, A., and Murshudov, G. N. (2018) Overview of refinement procedures within REFMAC5: utilizing data from different sources. Acta Crystallogr D Struct Biol 74, 215–227

61. Emsley, P., Lohkamp, B., Scott, W. G., and Cowtan, K. (2010) Features and development of Coot. Acta Crystallogr D Biol Crystallogr 66, 486–501

62. Chen, V. B., Arendall, W. B., 3rd, Headd, J. J., Keedy, D. A., Immormino, R. M., Kapral, G. J., Murray, L. W., Richardson, J. S., and Richardson, D. C. (2010) MolProbity: all-atom structure validation for macromolecular crystallography. Acta Crystallogr D Biol Crystallogr 66, 12–21

63. Shen, Y., Boulos, S., Sumrall, E., Gerber, B., Julian-Rodero, A., Eugster, M. R., Fieseler, L., Nystrom, L., Ebert, M. O., and Loessner, M. J. (2017) Structural and functional diversity in Listeria cell wall teichoic acids. J Biol Chem 292, 17832–17844

64. Wörmann, M. E., Corrigan, R. M., Simpson, P. J., Matthews, S. J., and Gründling, A. (2011) Enzymatic activities and functional interdependencies of Bacillus subtilis lipoteichoic acid synthesis enzymes. Mol Microbiol 79, 566–583

