## Supporting Information for "*Bacillus subtilis* YngB contributes to wall teichoic acid glucosylation and glycolipid formation during anaerobic growth"

List of Material included:
Supplemental Table 1
Supplemental Table 2
Supplemental Table 3
Supplemental Figure S1
Supplemental Figure S2
Supplemental Figure S3

**Supplemental Tables**

**Table S1. Observed and predicted masses of parental and MS/MS fragments for the glycolipids (32:0) [DAG-$Glc_2$ + $Na^+$] and (32:0) [DAG-$Glc_3$ + $Na^+$].**

| Observed mass | Predicted mass | Chemical formula |
|---|---|---|
| **(32:0) [DAG-$Glc_2$ + $Na^+$]** | | |
| 915.6 | 915.6 | (32:0) DAG-Glc-Glc + $Na^+$ |
| 673.5 | 673.3 | (17:0) MAG-Glc-Glc – $H_2O$ + $Na^+$ |
| 645.5 | 645.3 | (15:0) MAG-Glc-Glc –$H_2O$ + $Na^+$ |
| 405.2 | 405.1 | $CH_2$=CH-$CH_2$-Glc-Glc + $Na^+$ |
| 365.1 | 365.1 | Glc-Glc + $Na^+$ |
| 347.2 | 347.1 | Glc-Glc – $H_2O$ + $Na^+$ |
| 327.3 | 327.3 | (17:0) MAG – OH |
| 299.3 | 299.3 | (15:0) MAG – OH |
| **(32:0) [DAG-$Glc_3$ + $Na^+$]** | | |
| 1077.8 | 1077.7 | (32:0) DAG-Glc-Glc-Glc + $Na^+$ |
| 835.4 | 835.4 | (17:0) MAG-Glc-Glc-Glc – $H_2O$ + $Na^+$ |
| 807.4 | 807.4 | (15:0) MAG-Glc-Glc-Glc – $H_2O$ + $Na^+$ |
| 567.2 | 567.2 | $CH_2$=CH-$CH_2$-Glc-Glc-Glc + $Na^+$ |
| 527.2 | 527.2 | Glc-Glc-Glc + $Na^+$ |
| 509.1 | 509.2 | Glc-Glc-Glc – $H_2O$ + $Na^+$ |

MAG – monoacylgycerol; DAG – diacylglycerol; Glc – glucose. Note that only the observed masses for the glycolipids isolated from the wild-type strain are listed.

**Table S2. Bacterial strains used in this study**

| Unique ID | Strain name and resistance | Source |
|---|---|---|
| ***Escherichia coli* strains** | | |
| ANG127 | XL1-Blue | Stratagene |
| ANG191 | BL21(DE3) | (1) |
| ANG5206 | XL1-Blue pET28b-*gtaB*-cHis; KanR | This study |
| ANG5207 | XL1-Blue pET28b-*yngB*-cHis; KanR | This study |
| ANG5208 | BL21(DE3) pET28b-*gtaB*-cHis; KanR | This study |
| ANG5209 | BL21(DE3) pET28b-*yngB*-cHis; KanR | This study |
| ANG680 | DH5α pDR111; AmpR | (2) |
| ***Bacillus subtilis* strains** | | |
| ANG1691 | 168, trpC2 | (3) |
| ANG5277 | 168Δ*gtaB::kan*; KanR | (4) |
| ANG5263 | 168Δ*yngB::kan*; KanR | (4) |
| ANG5658 | 168Δ*gtaB::erm*; ErmR | (4) |
| ANG5659 | 168Δ*yngB::erm*; ErmR | (4) |
| ANG5675 | 168, *amyE::spec* $P_{hyper}$; SpecR (**Short: WT $P_{hyper}$**) | This study |
| ANG5676 | 168Δ*gtaB::kan amyE::spec* $P_{hyper}$; KanR SpecR (**Short: Δ*gtaB* $P_{hyper}$**) | This study |
| ANG5677 | 168Δ*gtaB::kan amyE::spec* $P_{hyper}$-*gtaB*; KanR SpecR (**Short: Δ*gtaB* $P_{hyper}$-*gtaB***) | This study |
| ANG5678 | 168Δ*gtaB::kan amyE::spec* $P_{hyper}$-*yngB*; KanR SpecR (**Short: Δ*gtaB* $P_{hyper}$-*yngB***) | This study |
| ANG5679 | 168Δ*yngB::kan amyE::spec* $P_{hyper}$; KanR SpecR (**Short: Δ*yngB* $P_{hyper}$**) | This study |
| ANG5680 | 168Δ*yngB::kan amyE::spec* $P_{hyper}$-*yngB*; KanR SpecR (**Short: Δ*yngB* $P_{hyper}$-*yngB***) | This study |
| ANG5681 | 168Δ*gtaB::erm* Δ*yngB::kan amyE::spec* $P_{hyper}$; ErmR KanR SpecR (**Short: Δ*gtaB*/Δ*yngB* $P_{hyper}$**) | This study |
| ANG5682 | 168Δ*gtaB::kan* Δ*yngB::erm amyE::spec* $P_{hyper}$-*gtaB*; ErmR KanR SpecR (**Short: Δ*gtaB*/Δ*yngB* $P_{hyper}$-*gtaB***) | This study |
| ANG5683 | 168Δ*gtaB::erm* Δ*yngB::kan amyE::spec* $P_{hyper}$-*yngB*; ErmR KanR SpecR (**Short: Δ*gtaB*/Δ*yngB* $P_{hyper}$-*yngB***) | This study |

**Table S3. Primers used in this study**

| Number | Name | Sequence |
|---|---|---|
| ANG3161 | 5-NcoI-pET28b-*gtaB*-cHis | CATGCCATGGGCAAAAAAGTACGTAAAGCCATAAT TCCAG |
| ANG3162 | 3-XhoI-pET28b-*gtaB*-cHis | CCGCTCGAGGCCGCTGCTGCCGCGCGGCACCAG GATTTCTTCTTTGTTTAGTAAACCTTC C |
| ANG3163 | 5-NcoI-pET28b-*yngB*-cHis | CATGCCATGGGCAGAAAAAAAGTGAGAAAAGCGGT TATAC |
| ANG3164 | 3-XhoI-pET28b-*yngB*-cHis | CCGCTCGAGGCCGCTGCTGCCGCGCGGCACCAG CCGCAGCATTTCTTTCGTTTCCCGTTTG |
| ANG111 | 5-pET28b-T7 promoter | TAATACGACTCACTATAGGG |
| ANG112 | 3-pET28b-T7 terminator | GCTAGTTATTGCTCAGCGG |
| ANG3197 | 5-*gtaB::kan* | TCATTTTAAATCATTTCATTCTTGATTC |
| ANG3198 | 3-*gtaB::kan* | ATTTATGTTTTCATTTAGTTTGTTTAAC |
| ANG3199 | 5-*yngB::kan* | TATTGCCGCGTCAGGGATCACCTT TTTG |
| ANG3200 | 3-*yngB::kan* | CTAATACAATCTCACTCGGAATGATTTC |
| ANG3203 | 5-HindIII-pDR111-*gtaB* | CCCAAGCTTAAAATAAGGAGGACCTTTTAAATGAAA AAAGTACGTAAAGC |
| ANG3204 | 3-NheI-pDR111-*gtaB* | CTAGCTAGCTTAGATTTCTTCTTTGTTTAGTAAACCT TC |
| ANG3205 | 5-HindIII-pDR111-*yngB* | CCCAAGCTTAAAATAAGGAGGAGAAGATGAATGAG AAAAAAAGTGAGAAAAG |
| ANG3206 | 3-NheI-pDR111-*yngB* | CTAGCTAGCTTACCGCAGCATTTCTTTCGTTTCCCGT TTG |
| ANG1663 | 5-pDR111-*amyE*-check | GCGAGGGAAGCGTTCACAGTTTCGGGC |
| ANG1664 | 3-pDR111-*amyE*-check | CGGTTGTAGCCCAAACGCCTTTCCGTGG |
| ANG1671 | 5-pDR111-seq | CGCTCTCCTGAGTAGGACAAATCCGC |
| ANG1672 | 3-pDR111-seq | CGGGAAACGGTCTGATAAGAGACACC |

**Supplemental Figures**

```
GtaB (B. subtilis)         --------MKKVRKAIIPAAGLGTRFLPATKAMPKEMLPIVDKPTIQYIIEEAVEAGIED 52
YngB (B. subtilis)         -------MRKKVRKAVIPAAGLGTRFLPATKAQPKEMLPIVDKPAIQYIVEEAAESGIED 53
YtdA (B. subtilis)         ----------MIKKAIIPAGGFGTRNLPVTKVIPKEMFPVGSKPVIHYLVEELKESGIED 50
A4JT02 (B. vietnamiensis)     -----MLKVTKAVFPVAGLGTRFLPATKASPKEMLPVVDKPLIQYAVEEAIEAGITE 52
GalU (C. glutamicum)       MSLPIDEHVNAVKTVVVPAAGLGTRFLPATKTVPKELLPVVDTPGIELIAAEAAELGATR 60
GalU (H. pylori)           ----------MIKKCLFPAAGYGTRFLPITKTIPKEMLPIVDKPLIQYAVEEAMEAGCEV 50
                                      : . :.*..* *** ** **. ***::*: ..* *.    *  * *

GtaB (B. subtilis)         IIIVTGKSKRAIEDHFDYSPELERNLEEKGKTELLEKVKKASN-LADIHYIRQKEPKGLG 111
YngB (B. subtilis)         ILIITGRNKRSIEDHFDRSAELEFNLREKGKTETLKEMQQIAD-LANIHYIRQKEPLGLG 112
YtdA (B. subtilis)         ILMVVSSHKNLIVDYFDSSLALEAFLASKNKLHLLR--EHPIP-DIRIHYVRQPYAKGLG 107
A4JT02 (B. vietnamiensis)  MIFVTGRSKRAIEDHFDKSYEIEAELEARGKEKLLSLVRSIKPSHVDCFYVRQAEALGLG 112
GalU (C. glutamicum)       LAIITAPNKAGVLAHFERSSELEETLMERGKTDQVEIIRRAAD-LIKAVPVTQDKPLGLG 119
GalU (H. pylori)           MAIVTGRNKRSLEDYFDTSYEIEHQIQGTNKENALKSIRNIIE-KCCFSYVRQKQMKGLG 109
                           : ::..  *  :  :*: *  :*  :   .* . :   .           : *    ***

GtaB (B. subtilis)         HAVWCARNFIGDEP--FAVLLGDDIVQAETPG--LRQLMDEYEKTLSSIIGVQQVPEEET 167
YngB (B. subtilis)         HAVLCAEHFIGDEP--FAVLLGDDIMVSETPA--LRQLMDVYDVYGTEVVGVQSVLPEDV 168
YtdA (B. subtilis)         DAISFGKQFAGGEP--FAVVLPDDLIFSANQP-ALGQLIEAYTKYQSSVIGLKETKTEDL 164
A4JT02 (B. vietnamiensis)  HAVLCAEKLVGDNP--FAVILADDLLDGPTPV--LRQMIDVFDHYHASVIGVEEIAPADS 168
GalU (C. glutamicum)       HAVGLAESVLDDDEDVVAVMLPDDLVLPTGVM---ERMAQVRAEFGGSVLCAVEVSEADV 176
GalU (H. pylori)           HAILTGEALIGNEP--FAVILADDLCISHDHPSVLKQMTSLYQKYQCSIVAIEEVALEEV 167
                           .*:  .. . ..:   .**:* **:           :: .       .::   .    :

GtaB (B. subtilis)         HRYGIIDPLTSEGR--RYQVKNFVEKPPKGTAPSNLAILGRYVFTPEIFMYLEEQQVGAG 225
YngB (B. subtilis)         SKYGIINTSGSQGH--VYEVNDLVEKPSPEEAPSEIAVMGRYVLNSSIFSVLKTIGRGAG 226
YtdA (B. subtilis)         HHYGVIKGEPVEKG--LYRIQDIVEKPKQ-NPPSHFAAAGRYIFTPDIFNELEALEADSG 221
A4JT02 (B. vietnamiensis)  KSYGVIDGKRWEDD--LFKLSGIVEKPEPAQAPSNFGVVGRYVLKPKIFKHLRGLKPGAG 226
GalU (C. glutamicum)       SKYGIFEIEADTKDSDVKKVKGMVEKPAIEDAPSRLAATGRYLLDRKIFDALRRITPGAG 236
GalU (H. pylori)           SKYGVIRGEWLEEG--VYEIKDMVEKPNQEDAPSNLAVIGRYILTPDIFEILSETKPGKN 225
                             **::            .:..:****    .**.:.  ***::  .**  *     . .

GtaB (B. subtilis)         GEIQLTDAIQKLNEI-QRVFAYDFEGKRYDVGEKLGFITTTLEFAMQDKELRDQLVPFME 284
YngB (B. subtilis)         NEIQLTDALREVCRK-EPIHARLLEGNRYDIGDKLGCFKASTEIGLMRPEMRSQLLAYLE 285
YtdA (B. subtilis)         GEVQVTDAIKASLGA-CTVYGKLLEGERYDIGLQKDYLKLIYDMLKTEKNPQ-------- 272
A4JT02 (B. vietnamiensis)  GELQLTDAIQSLLTD-EQVLAYRYDGTRFDCGSKLGYLKATVEFALRHPEVAADFERYLL 285
GalU (C. glutamicum)       GELQLTDAIDLLIDEGHPVHIVIHQGKRHDLGNPGGYIPACVDFGLSHPVYGAQLKDAIK 296
GalU (H. pylori)           NEIQITDALRTQAKR-KRIIAYQFKGKRYDCGSVEGYIEASNAYYKKRLLE--------- 275
                           .*:*:***:         :     .* *.* *   . :

GtaB (B. subtilis)         GLLNKEEI------------- 292
YngB (B. subtilis)         DVIKRETKEMLR--------- 297
YtdA (B. subtilis)         ---------------------
A4JT02 (B. vietnamiensis)  ARMSEQLVV------------ 294
GalU  (C. glutamicum)      QILAEHEAAERIADDSQVKLE 317
GalU  (H. pylori)          ---------------------
```

**Figure S1. Amino acid sequence alignment of *B. subtilis* UGPases and UGPases with available UDP-glucose bound structures.** The amino acid sequences of the *Bacillus subtilis* 168 confirmed and predicted UGPases GtaB, YngB and YtdA were aligned with the protein sequence of A4JT02 from *Bulkholderia vietnamiensis,* GalU from *Corynebacterium glutamicum* and GalU from *Helicobacter pylori,* for which UDP-glucose bound structures are available using CLUSTALW (5). Based on the alignment and the structural information, possible UDP-glucose binding residues were identified and highlighted in yellow and a conserved aspartic acid likely involved in metal ion binding is highlighted in cyan.

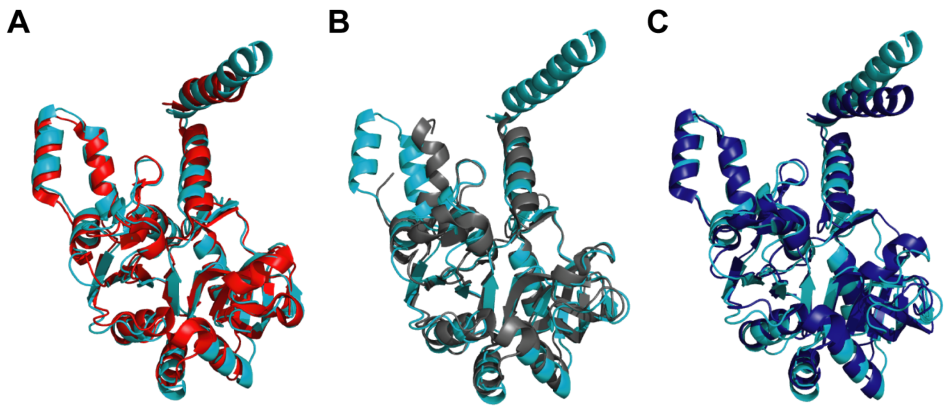


**Figure S2. Superimposition of the *B. subtilis* YngB structure with structures of other UGPases.** *A,* Superimposition of the *B. subtilis* YngB structure (cyan in all panels) with A4JT02 (PDB code: 5i1f) from *Bulkholderia vietnamiensis* (red) yielding an rmsd value of 1.39Å over 274 aligned residues*;* *B*, superimposition with $GalU_{Hp}$ (PDB code: 3juk) from *Helicobacter pylori* (grey) giving an rmsd value of 1.61 Å over 250 aligned residues; and *C*, superimposition with $GalU_{Cg}$ (PDB code 2pa4) from *Corynebacterium glutamicum* (blue) yielding an rmsd value of 2.11 Å over 282 aligned residues.

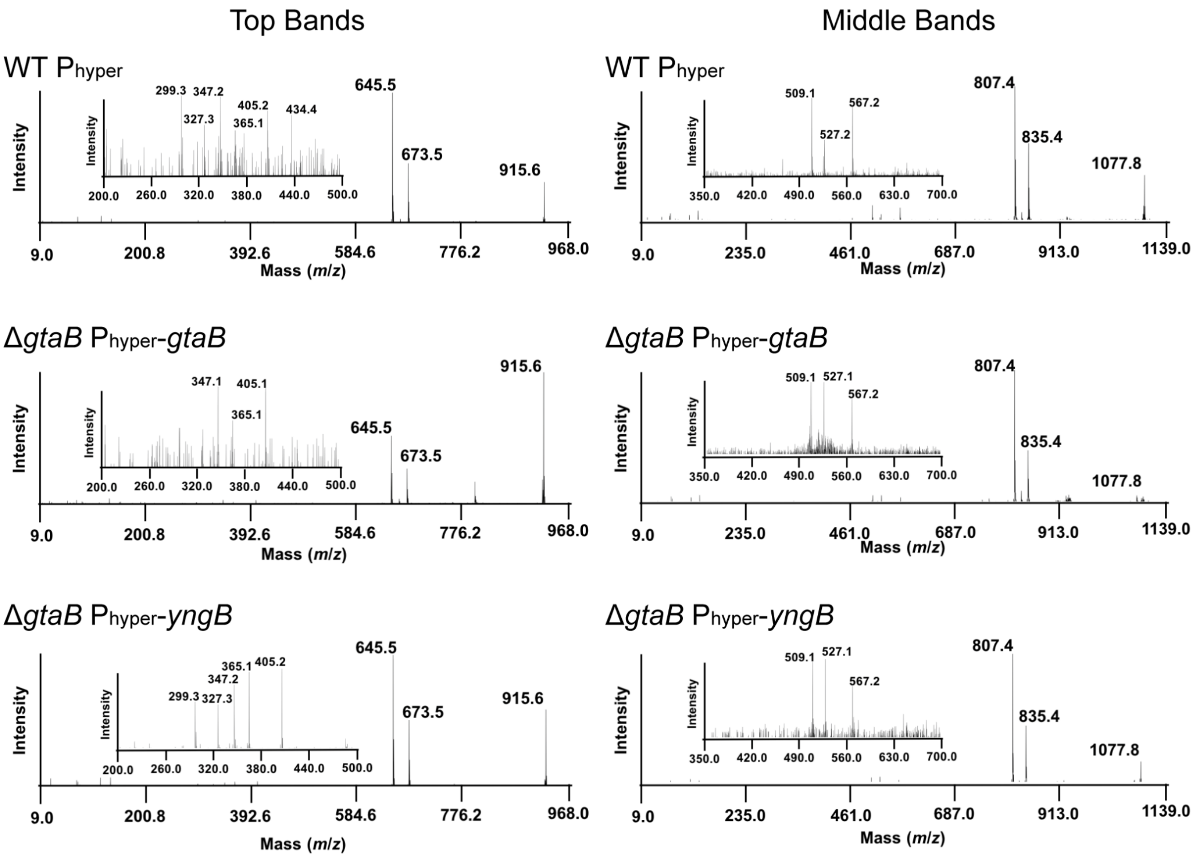


**Figure S3. MALDI TOF MS/MS analysis of extracted glucolipids.** Tandem mass spectrometry was performed on glycolipids isolated from one of the experiments from the wild-type *B. subtilis* (WT $P_{hyper}$) strain and the *gtaB* complementation strains Δ*gtaB* $P_{hyper}$-*gtaB* and Δ*gtaB* $P_{hyper}$-*yngB*. The analysis was performed on the glycolipids with an observed mass of 915.6 (*m*/*z*) (top band) and proposed to correspond to (32:0) [DAG-$Glc_2$ + $Na^+$] and an observed mass of 1077.9 (*m*/*z*) (middle band) and proposed to correspond to (32:0) [DAG-$Glc_3$ + $Na^+$]. The MS/MS spectra are shown in the figure and chemical formula for the fragmented species are listed in Table S1. The insert is an enlarged view of the species in the 200-550 *m*/z range for the glycolipids isolated from the top band or 350-700 *m*/*z* range for the glycolipids isolated from the middle bands.

### References


1. Studier, F. W., Rosenberg, A. H., Dunn, J. J., and Dubendorff, J. W. (1990) Use of T7 RNA polymerase to direct expression of cloned genes. *Methods Enzymol* **185**, 60-89
2. Wagner, J. K., Marquis, K. A., and Rudner, D. Z. (2009) SirA enforces diploidy by inhibiting the replication initiator DnaA during spore formation in *Bacillus subtilis*. *Mol Microbiol* **73**, 963-974
3. Burkholder, P. R., and Giles, N. H., Jr. (1947) Induced biochemical mutations in *Bacillus subtilis*. *Am J Bot* **34**, 345-348
4. Koo, B. M., Kritikos, G., Farelli, J. D., Todor, H., Tong, K., Kimsey, H., Wapinski, I., Galardini, M., Cabal, A., Peters, J. M., Hachmann, A. B., Rudner, D. Z., Allen, K. N., Typas, A., and Gross, C. A. (2017) Construction and Analysis of Two Genome-Scale Deletion Libraries for *Bacillus subtilis*. *Cell Syst* **4**, 291-305 e297
5. Madeira, F., Park, Y. M., Lee, J., Buso, N., Gur, T., Madhusoodanan, N., Basutkar, P., Tivey, A. R. N., Potter, S. C., Finn, R. D., and Lopez, R. (2019) The EMBL-EBI search and sequence analysis tools APIs in 2019. *Nucleic Acids Res* **47**, W636-W641
